# Deep Learning for RNA Synthetic Biology

**DOI:** 10.1101/872077

**Authors:** Nicolaas M. Angenent-Mari, Alexander S. Garruss, Luis R. Soenksen, George Church, James J. Collins

## Abstract

Engineered RNA elements are programmable tools capable of detecting small molecules, proteins, and nucleic acids. Predicting the behavior of these tools remains a challenge, a situation that could be addressed through enhanced pattern recognition from deep learning. Thus, we investigate Deep Neural Networks (DNN) to predict toehold switch function as a canonical riboswitch model in synthetic biology. To facilitate DNN training, we synthesized and characterized *in vivo* a dataset of 91,534 toehold switches spanning 23 viral genomes and 906 human transcription factors. DNNs trained on nucleotide sequences outperformed (R^2^=0.43-0.70) previous state-of-the-art thermodynamic and kinetic models (R^2^=0.04-0.15) and allowed for human-understandable attention-visualizations (VIS4Map) to identify success and failure modes. This deep learning approach constitutes a major step forward in engineering and understanding of RNA synthetic biology.

**One Sentence Summary:** Deep neural networks are used to improve functionality prediction and provide insights on toehold switches as a model for RNA synthetic biology tools.

## Main Text

Engineered ribonucleic acid (RNA) molecules with targeted biological functions play an important role in synthetic biology (*1*), particularly as programmable response elements for small molecules, proteins, and nucleic acids. Examples include riboswitches, riboregulators, and ribozymes, many of which hold great promise for a variety of *in vitro* and *in vivo* applications (*1, 2*). Despite their appeal, the design and validation of this emerging class of synthetic biology modules have proven challenging due to variability in function that remains difficult to predict (*2–9*). Current efforts aiming to unveil fundamental relationships between RNA sequence, structure, and behavior focus mostly on mechanistic thermodynamic modeling and low-throughput experimentation, which often fail to deliver sufficiently predictive and actionable information to aid in the design of complex RNA tools (*2–9*). Deep learning, by contrast, constitutes a set of computational techniques well suited for feature recognition in complex and highly combinatorial biological problems (*10–14*), such as the sequence design space of synthetic RNA tools. However, the application of deep learning to predicting function in RNA synthetic biology has been limited by a notable scarcity of datasets large enough to effectively train deep neural networks. Toehold switches, in particular, represent a benchmark RNA element in synthetic biology that could greatly benefit from deep learning approaches to better predict function and elucidate useful design rules.

Toehold switches are a class of versatile prokaryotic riboregulators inducible by the presence of a fully programmable trans-RNA trigger sequence (*2–6, 15, 16*). These RNA synthetic biology modules have displayed impressive dynamic range and orthogonality when used both *in vivo* as genetic circuit components (*2, 5, 6*), and *in vitro* as nucleic acid diagnostic tools using cell-free protein synthesis (CFPS) systems (*3, 4, 15, 16*). Similar to other RNA synthetic biology tools, a substantial fraction of toehold switches show poor to no measurable function when tested experimentally, and while efforts have been made to establish rational, mechanistic rules for improved performance based on low-throughput datasets (*2–9, 15, 16*), the practical utility of these approaches remains inconclusive. Thus, considering the wide applicability and general challenges of toehold switch design, our objective in this study was to develop a deep learning platform to predict toehold switch function as a canonical RNA switch model in synthetic biology.

To achieve our goal, we first aimed to expand the size of available toehold datasets using a high-throughput DNA synthesis and sequencing pipeline to characterize over 10^5^ new toehold switches. We then used this comprehensive new dataset to demonstrate that deep neural networks trained directly on switch RNA sequences can outperform rational thermodynamic and kinetic analyses to predict toehold switch function. Furthermore, we enhanced the transparency of our deep learning approach by utilizing a nucleotide complementarity matrix input representation to visualize important learned secondary structure patterns in selected models. This attention-visualization technique, which we term VIS4Map (<u>Vis</u>ualizing <u>S</u>econdary <u>S</u>tructure <u>S</u>aliency <u>Map</u>s), allowed us to identify RNA module success and failure modes by discovering secondary structures that our deep learning model used to accurately predict toehold switch function. The resulting dataset, models, and visualization analysis (Fig. 1) represent a substantial step forward for the validation and interpretability of high-throughput approaches to designing RNA synthetic biology tools, surpassing the limits of current mechanistic RNA secondary structure modeling.

**Fig.1.**
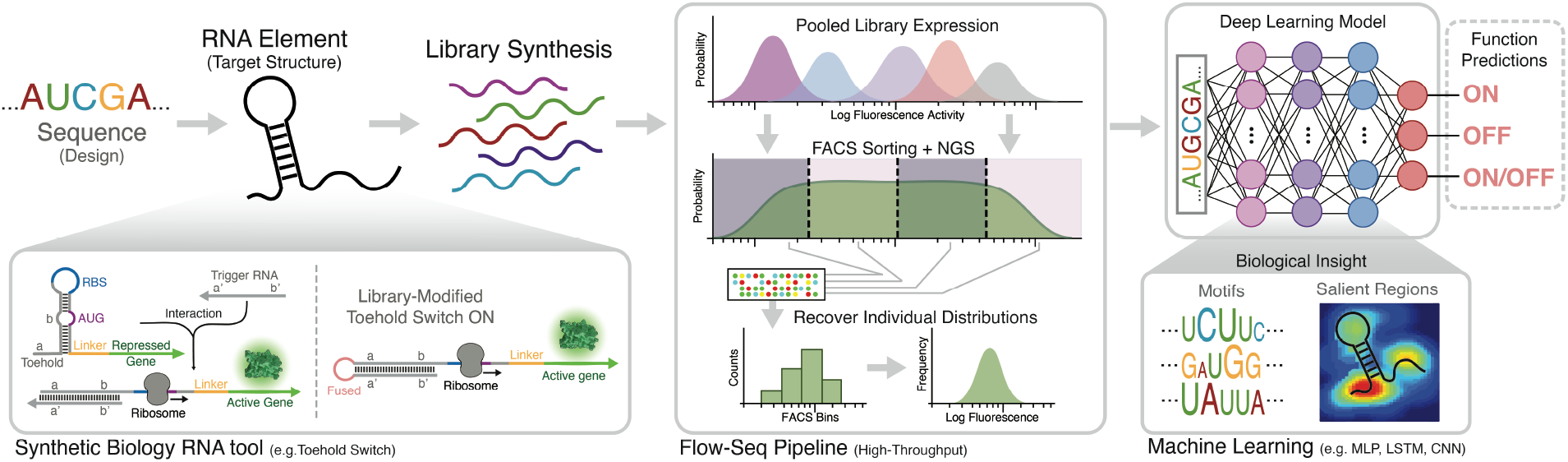
Deep learning for RNA synthetic biology pipeline. RNA tool selection is followed by library synthesis and characterization with analysis using deep neural networks (DNN) to provide functionality predictions and design insight. We used a high-throughput toehold switch library as a canonical model for the general investigation of RNA synthetic biology tools. The original toehold switch architecture from Green et al. (*2*) was used, containing a 12-nucleotide toehold (a/a’) and an 18-nucleotide stem (b/b’) fully unwound by the trigger (left-bottom). We selected to fuse the RNA trigger to the 5’ end of the switch by an unstructured linker to facilitate library synthesis. Then, a flow-sequence (seq) pipeline was used to characterize the fluorescence signal of individual toehold switches in a pooled sequential assay, including pooled induction, FACS sorting, next-generation sequencing (NGS) and count frequency analysis. Finally, various DNN architectures were used to predict data outputs, while features contributing to DNN predictions were intuitively visualized to elucidate biological insights.

### Library synthesis, characterization, and validation

A fundamental hurdle in applying deep learning techniques to RNA synthetic biology systems is the limited size of currently published datasets, which are notably smaller than typical dataset sizes required for the training of deep network architectures in other fields (*10, 17–21*). For example, to date, less than 1000 total toehold switches have been designed and tested (*2–6, 9, 15, 16*), a situation that currently limits the synthetic biology community’s ability to analyze this type of response molecule using deep learning techniques. Towards improving our understanding and ability to predict new functional RNA-based response elements, we synthesized and characterized an extensive *in vivo* library of toehold switches using a high-throughput flow-seq pipeline (*22*) for subsequent exploration using various machine learning and deep learning architectures.

Our toehold switch library was designed and synthesized based on a large collection (244,000) of putative trigger sequences, spanning the complete genomes of 23 pathogenic viruses, the entire coding regions of 906 human transcription factors, and ~10,000 random sequences. From a synthesized oligo pool, we generated two construct libraries, for ON and OFF states, which were subsequently transformed into BL21 *Escherichia coli* (Fig. 1, S1A,B). The first library contained OFF toehold switch constructs that lacked a trigger, while the second library of ON constructs contained the same toeholds with complementary triggers fused to their corresponding switches. The two libraries were then sorted on a fluorescence-activated cell sorter (FACS) using four bins (Fig. 1, S2), and the toehold switch variants contained in each bin were quantified using next-generation sequencing (NGS) to recover their individual fluorescence distributions from raw read counts (Fig. 1). After quality control (Table S1), the toehold switch library contained 109,067 ON state measurements (Fig. 2A), 163,967 OFF state measurements (Fig. 2B), and 91,534 ON/OFF paired ratios (Fig. 2C), where both ON and OFF states were characterized for a given switch (Fig. 2E,F). ON and OFF data were normalized from 0 to 1, resulting in an ON/OFF ratio normalized from −1 to 1 (see Supplementary methods section).

**Fig.2.**
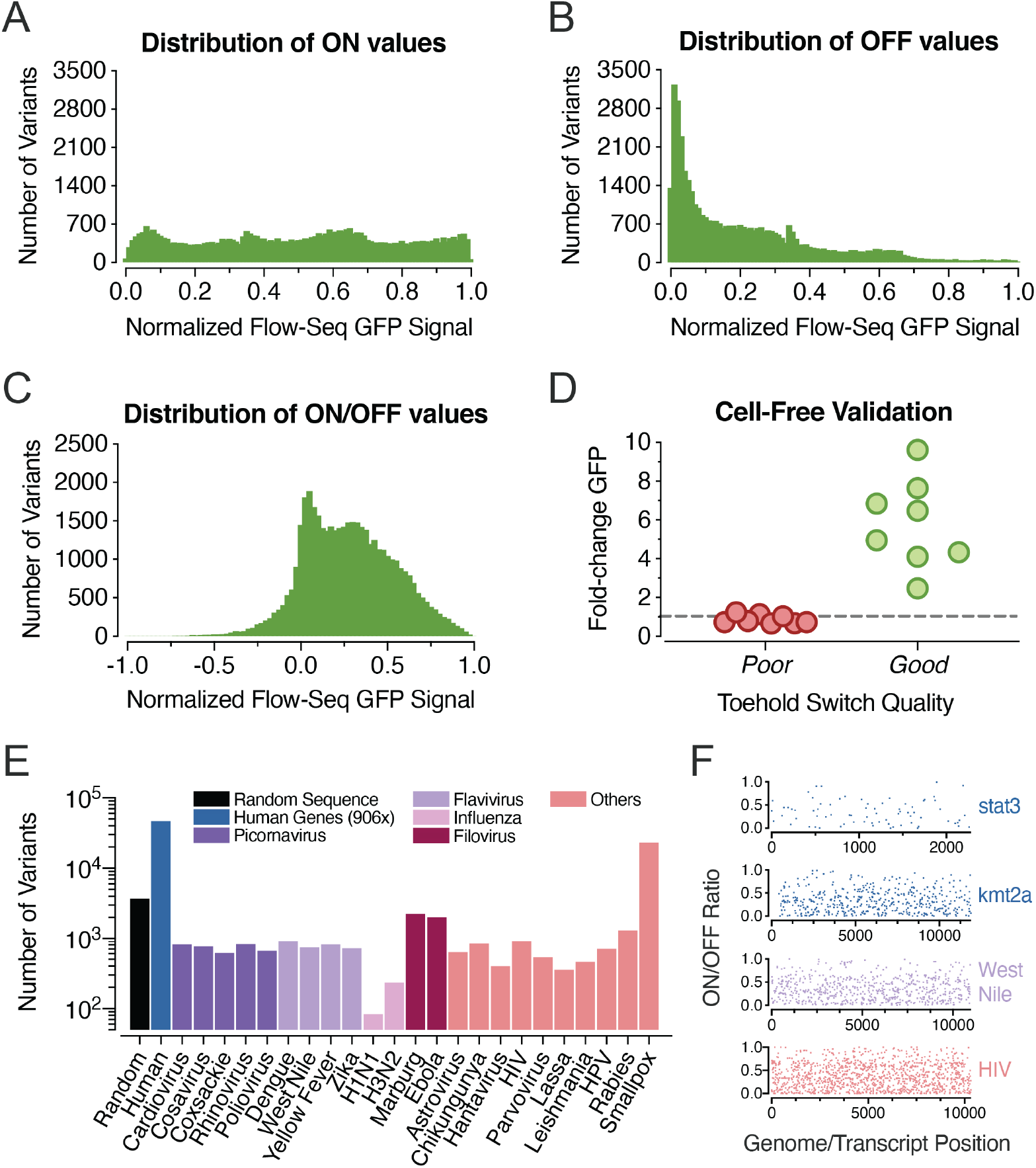
Flow-seq toehold switch library characterization and trigger ontology. The distribution of recovered toeholds for (A) ON-state signals, (B) OFF-state signals, and (C) calculated ON/OFF ratios are shown (selected from quality control process #3, QC3 in Fig. S13 and Table S1). (D) Validation results for toehold switches expressed in a PURExpress cell-free system with un-fused trigger RNA, including eight low-performing (poor, ON/OFF<0.05) and eight high-performing (good, ON/OFF>0.97) samples. Obtained *in vivo* flow-seq data show competency in classifying switch performance for this *in vitro* cell-free biological context. (E) Tested switch/trigger variants from each origin category, including randomly generated sequences, 906 human transcription factor transcripts, and 23 pathogenic viral genomes. (F) Experimental ON/OFF ratios for all triggers tiled across the transcripts of two clinically relevant human transcription factors (*stat3* and *kmt2a*) upregulated in cancerous phenotypes (*42, 43*), as well as all triggers tiled across the genomes of two pathogenic viruses: West Nile Virus (WNV) and Human Immunodeficiency Virus (HIV). GFP= Green Fluorescent Protein; Seq=Sequence; HPV=Human Papillomavirus.

Since RNA synthetic biology tools such as toehold switches are often used within *in vitro* cell-free systems (*3, 4, 15, 16*), we validated our *in vivo* ON/OFF measurements in an *in vitro* setting to ensure they were reasonable indicators of switch performance in a CFPS system. To achieve this, we selected eight high-performance switches and eight low-performance switches, and individually cloned and characterized them in a PURExpress CFPS (Fig. 1D, S5 & Table S2). All low-performance switches showed no induction, while the high-performance switches showed a spread of ON/OFF ratios between 2 and 10 (p<0.0001 between high and low switches, two-tailed t-test). These results confirm that while the performance of toehold switches *in vivo* and *in vitro* may differ, *in vivo* measurements can still be used to classify categorically whether a switch will function *in vitro*.

### Rational analysis using thermodynamic RNA secondary structure models

Before initiating the exploration of deep learning models to predict function in our large-scale toehold switch library, we sought to determine whether traditional tools for analyzing synthetic RNA modules could be used to accurately predict toehold switch behavior, including k-mer searches and mechanistic modeling using thermodynamic and kinetic parameters. K-mer searches of biological sequence data are often used to discover motifs, and while certain overrepresented motifs were found in our dataset (Fig. 3A & Table S3), utilization of these did not significantly improve functional predictions of switch behavior. Other current state-of-the-art approaches for designing RNA synthetic biology tools primarily analyze secondary structure using thermodynamic principles (*23–25*). Following such prior works, we used NUPACK (*23*) and ViennaRNA (*25*) software packages to calculate a total of 30 rational features for our entire library, including the minimum free energy (MFE), ideal ensemble defect (IED), and native ensemble defect (NED) of the entire toehold switch library as well as various sub-segments in each sequence (Table S4). A number of these parameters had previously been reported to correlate with experimental toehold switch ON/OFF measurements for smaller datasets (*2*), and NUPACK’s design algorithm, in particular, is set to optimize IED when proposing target RNA secondary structures (*3, 4, 15, 23*). However, when analyzing these rational features with our larger dataset, we found them to be poor predictors of toehold switch function (Fig. 3B, S6). In modest agreement with the findings of Green et al. (*2*), the MFE of the RBS-linker region showed the highest correlation of this feature set for ON/OFF (R^2^: ON=0.14, OFF=0.06, ON/OFF=0.04), with NUPACK’s IED also showing above-average correlation (R^2^: ON=0.07, OFF=0.02, ON/OFF=0.03). While measurable, these correlation metrics were far too weak for practical use in computer-aided design of this specific RNA synthetic biology tool (*3, 4, 15, 23*).

**Figure 3.**
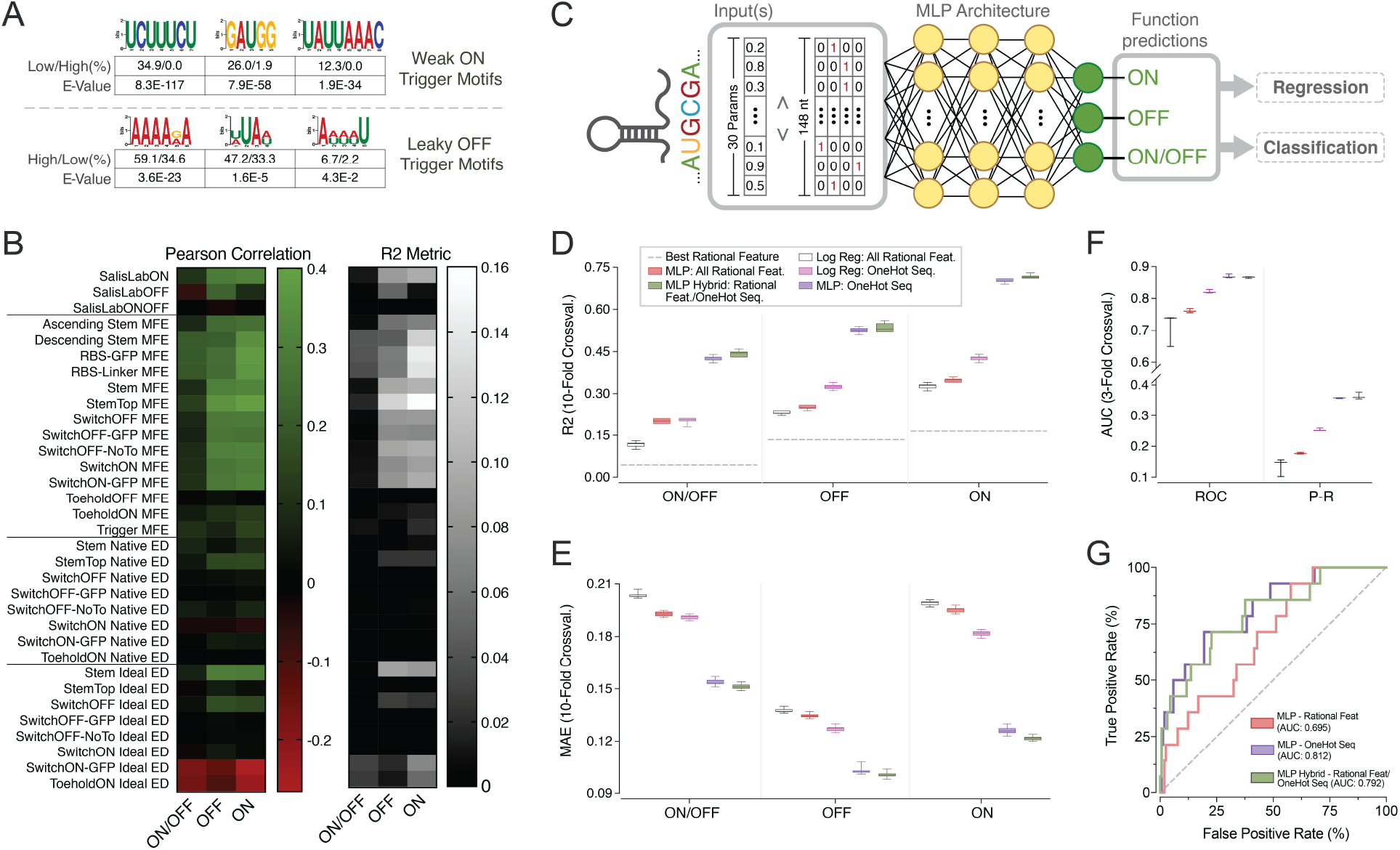
Analysis of toehold switch performance using sequence k-mers, rational thermodynamic features, and sequence-based multilayer perceptron (MLP) models. (A) Sequence logos for k-mer motifs discovered to be disproportionately represented in weakly induced switches (low ON) and leaky switches (high OFF), functional proportions, and E-values. (B) The Pearson correlation (left, |*max*|=0.4) and R^2^ metric (right, |*max*|=0.16) for thirty state-of-the-art thermodynamic features and obtained RBS Calculator v2.1 outputs. (C) Base architecture of investigated MLP models, featuring three fully connected layers. For training in regressionmode, three different outputs were predicted (ON, OFF, ON/OFF), whereas for classification training only a single binary output based on ON/OFF (threshold at 0.7) was predicted. (D) Box and whisker plots for R^2^ between experimental and regression-based predictions for best performing rational features, logistic regression models and MLPs. (E) Mean absolute error (MAE) between experimental and predicted values for these same models. (F) Box and whisker plots for area under the curve (AUC) of the receiver-operator curve (ROC) and the precisionrecall curve (P-R) in classification-mode predictions compared to experimental values. In both regression and classification, the one-hot encoded sequence MLP delivered top-in-class performance without using pre-computed thermodynamic or kinetic metrics. (G) ROC curves of pre-trained MLP classification models validated with an unseen 168-sequence external dataset from Green et al. (*2*).

We next explored the use of more complex thermodynamic models that take into account well-established hypotheses for translation initiation and the ribosome docking mechanism in combination with multiple thermodynamic features to improve their predictions (*26–31*). One of the most developed of these models is the Ribosome Binding Site (RBS) calculator (v2.1; Salis Lab); a comprehensive regression model parameterized on thousands of curated RBS variants (*26–29*). We used the RBS calculator to predict the ON and OFF translation initiation rates for our toehold switches, but also found low predictive performance comparable to other rational features (Fig. 3B) when tested on our database (R^2^: ON=0.09, OFF=0.05, ON/OFF=0.0001).

One potential explanation for the limited predictive power of current thermodynamic models for RNA folding tasks concerns the influence of kinetically stable secondary structure intermediates that may compete with thermodynamic equilibrium states (*29, 32*). To determine whether a kinetic analysis of toehold switch folding dynamics could help explain our experimental results, we calculated four additional features based on kinetic trajectories using the Kinfold package (*33*) (Fig. S7). As with predictions obtained using other thermodynamic models, these kinetic features showed poor correlations (R^2^: ON=0.04, OFF=0.04, ON/OFF=0.001 for the best feature) to our empirical dataset (Fig. S7E). Considering these results, the cause of limited functional predictions from thermodynamic and kinetic RNA secondary structure models remains unclear but may stem from the use of potentially incomplete energetic models, incorrect mechanistic hypotheses, or interference from the *in vivo* context of the bacterial cell. Regardless of the source of error, we sought to explore deep learning as a machine learning paradigm to develop models with higher predictive abilities than previously reported, with the hope of allowing useful computer-aided systems for the design of RNA synthetic biology tools.

### Improved prediction using sequence-based multilayer perceptron models

Given that simple regression models based on previous state-of-the-art RNA thermodynamic and kinetic calculations were ineffective at predicting toehold switch performance, we next tested the use of feed-forward neural networks, also known as multilayer perceptron (MLP) models, as a baseline architecture for our investigation (Fig. 3C). We first trained a three-layer MLP model on our dataset with an input consisting of the 30 previously calculated thermodynamic rational features (see Methods section for further detail). When trained in regression mode, this MLP model was able to deliver better predictions than any of the individual rational features or the RBS calculator based on R^2^ and mean absolute error (MAE) (R^2^: ON=0.35, OFF=0.25, ON/OFF=0.20) (Fig. 3D, E). Similarly, when this model was trained in classification mode (ON/OFF: binarized at +/-0.7), as seen in Fig. S8, it achieved a 0.76 area under the receiver-operator curve (AUROC) and 0.18 area under the precision-recall curve (AUPRC), as seen in Fig. 3F. The MLP model slightly outperformed a logistic regressor trained on the same rational features (Fig. 3D,E,F), suggesting that the MLP architecture was able to abstract higher-order patterns from these features as compared to simpler non-hierarchical models.

While these results already constitute an improvement compared to the current state-of-the-art analysis of RNA synthetic biology tools, we hypothesized that the use of pre-computed rational features as network input led to information loss that could inherently limit the predictive power of these models. Considering that possibility, we trained an MLP model solely on one-hot encoded sequence representations of our toehold switches, eliminating potential bias introduced by *a priori* mechanistic modeling. We found that this sequence-based MLP delivered improved functional predictions based on R^2^ and MAE metrics (R^2^: ON=0.70, OFF=0.53, ON/OFF=0.43) (Fig. 3D, E, S9). These values represent a doubling of R^2^ performance as compared to the MLP trained on rational features and a ten-fold improvement in ON/OFF R^2^ over the best individual rational feature used for previous linear models. When training for classification, our one-hot sequence MLP produced similarly improved AUROCs and AUPRCs of 0.87 and 0.36, respectively (Fig. 3F).

The improvement in performance when training on sequence-only inputs compared to rational features suggests that significant information loss occurs when performing thermodynamic calculations on toehold switch sequences, a problem that may extend to other RNA synthetic biology tools in use today. The sequence-only MLP model dramatically outperformed a logistic regressor model trained on the same one-hot sequence input (Fig. 3D,E,F), further supporting the hypothesis that improved accuracy of our sequence-based MLP arises from learned hierarchical non-linear features extracted directly from RNA sequences. Concatenating both the rational features and the one-hot representation into a combined input gave a small but significant improvement in regression mode (ΔR^2^ ≈ 0.025 and ΔMAE ≈ −0.0025, p<0.05 for all six comparisons, two-tailed t-test), but no significant improvement for AUROC or AUPRC when in classification mode (Fig. 3D,E,F). These results suggest that while the use of rational features may facilitate the abstraction of potentially relevant information of toehold switch function, the one-hot sequence-only MLP model can recover such information without *a priori* hypothesis-driven assumptions built into the model if given sufficient training data.

In order to evaluate the degree of biological generalization in our sequence-only MLP model, we performed two additional rounds of validation. First, we iteratively withheld each of the 23 tiled viral genomes in the dataset during training and predicted their function as test sets, resulting in a 0.82-0.98 AUROC range (average 0.87, Fig. S10), similar to previous results from our sequence-only MLP. We then carried out an external validation on unseen data from a previously published dataset of 168 characterized toehold switches (*2*) that had been collected under different experimental conditions. Our MLP models achieved an AUROC of 0.70, 0.81, and 0.79, when trained on rational features, one-hot sequence, and concatenated inputs, respectively (Fig. 3G). The improved performance observed when training the models directly on nucleotide sequence rather than thermodynamic features, even for an external dataset, suggest a competent degree of biological generalization and supports the value of modeling RNA synthetic biology tools using deep learning and high-throughput datasets, removing the current assumptions of mechanistic rational parameters.

### Predictive performance of higher-capacity deep learning models

Having explored a baseline deep learning architecture, we next sought to determine whether training our dataset on higher-capacity convolutional neural networks (CNN) and long shortterm memory (LSTM) recurrent neural networks could increase our predictive ability. CNN and LSTM models have been applied to a variety of biological datasets in recent years, and have been cited as being particularly adept at recognizing motifs and long-range interactions in nucleotide sequence data (*10, 17–20, 34–38*). We trained a CNN on a one-hot sequence input, an LSTM on a one-hot sequence input, and a CNN on a two-dimensional (2D), one-hot complementarity map representation input (see Methods for complete descriptions of all models). Upon evaluating both the R^2^ and MAE in regression mode and the AUROC and AUPRC in classification mode for these models (Fig. 4A,B,C,D), we concluded that these neural network architectures did not lead to superior predictive models, as compared to the sequencebased, three-layer MLP described previously. In these cases increased model capacity led to under-or over-fitting, requiring additional training examples or improved fine-tuning to accelerate effective training.

**Figure 4.**
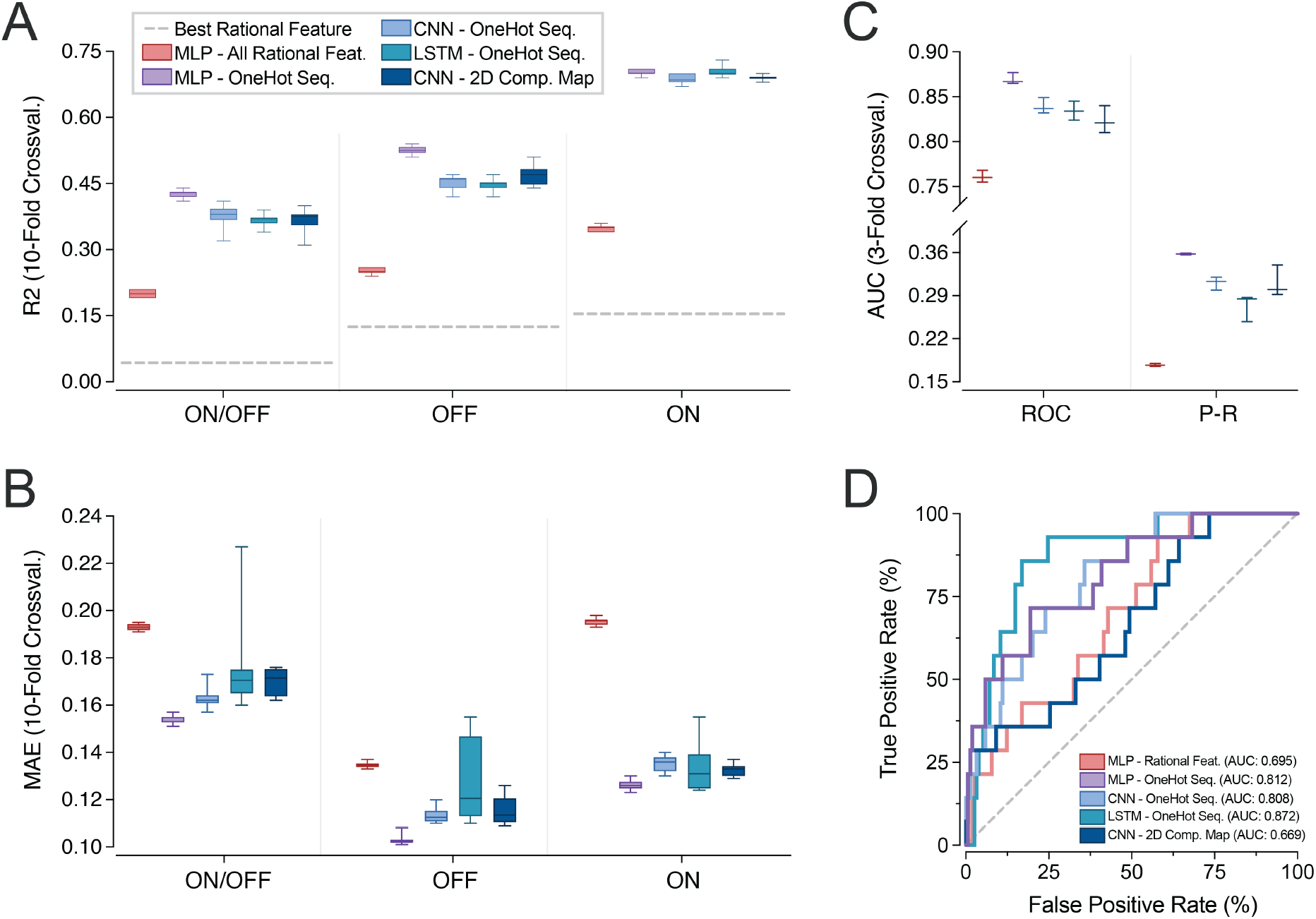
Evaluation of neural network architectures with increased capacity. Performance metrics for convolutional neural networks (CNN) and long short-term memory (LSTM) networks trained on one-hot encoded toehold sequences, as well as a CNN trained on a twodimensional, one-hot encoded sequence complementarity map. All models are compared to the previously reported MLPs trained on the 30 pre-calculated thermodynamic features and one-hot toehold sequences. For regression-based predictions (A) shows box and whisker plots for R^2^ metric, while (B) shows mean absolute error (MAE) for all models. In the case of classificationbased predictions (C) shows box and whisker plots of the area under the curve (AUC) of the receiver-operator curve (ROC) and the precision-recall curve (P-R) for all tested models. In both regression and classification, the one-hot encoded sequence MLP delivered top-in-class performance as compared to higher capacity deep learning models. (D) ROC curves of pretrained higher-capacity classification models validated with an unseen 168-sequence external dataset from Green et al. (*2*).

### Visualizing learned RNA secondary structure motifs with VIS4Map

One significant drawback of using deep learning to predict biological function is the inherent difficulty in understanding learned patterns in a way that helps researchers to elucidate biological mechanisms underlying model predictions. By contrast, mechanistic hypothesis-driven models can more directly inform which aspects of a biological theory best explain the observations. Various methods have been established to address this limitation, including alternative network architectures (*39*), and the use of saliency maps (*40, 41*), which reveal the regions of an input that deep learning models weigh most heavily and therefore pay the most attention to when making predictions. While saliency maps have been previously used to visualize model attention in one-hot representations of sequence data (*10, 17, 18, 20, 40*), such implementations focus only on the primary sequence and have not been developed to identify secondary structure interactions, which are especially relevant in the operation of RNA synthetic biology elements. In the few cases where secondary structure has been investigated, input representations have been constrained to predetermined structures based on the predictions of thermodynamic models (*37, 38*) whose abstractions we have found cause significant information loss.

We sought to visualize RNA secondary structures learned by our neural networks in a manner unconstrained by thermodynamic modeling. To achieve this, we trained a CNN on a twodimensional nucleotide complementarity map representation (Fig. 5A) to allow for attention pattern visualization in this secondary structure space. Each position in this complementarity map corresponds to the potential pair between two nucleotides, indicating its identity with a one-hot encoding (G-C, C-G, A-U, U-A, G-U, U-G, or an unproductive pair). We hypothesized that by training deep networks on such a representation of RNA sequences, it would be possible for generated saliency maps to reveal learned secondary structures as visually intuitive diagonal features. Importantly, because the complementarity map is unconstrained by *a priori* hypotheses of RNA folding (similarly to our sequence-based MLP models), we anticipated this approach to be able to identify secondary structures that might be overlooked by commonly used thermodynamic and kinetic algorithms, such as NUPACK and Kinfold.

**Figure 5.**
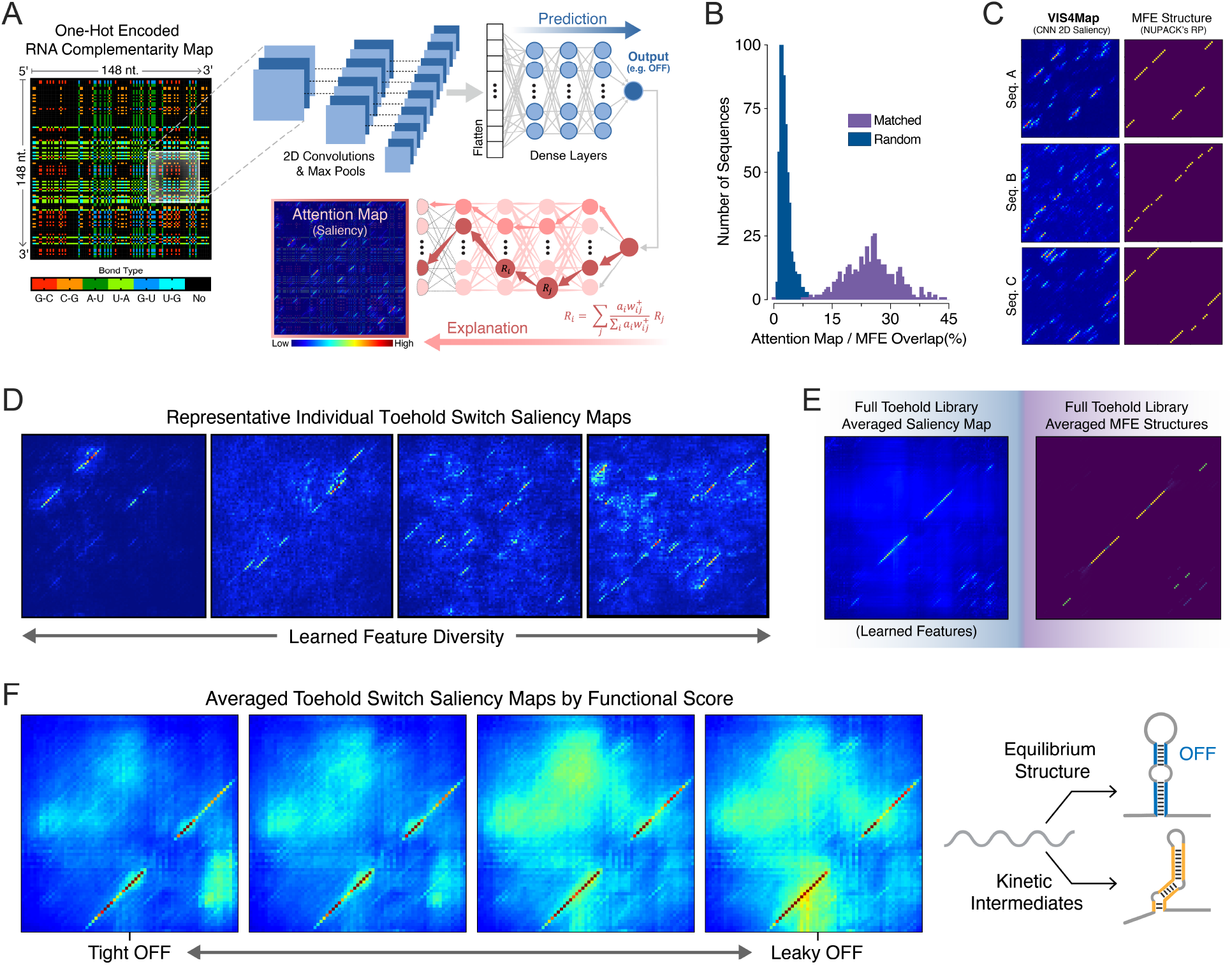
VIS4Map: Visualizing secondary structure features using saliency maps of a sequence-based complementarity matrix input. (A) A simplified schematic of the CNN-based architecture used to generate toehold functional predictions with network attention visualizations. The system receives a one-hot encoded, two-dimensional (2D) sequence complementarity map as input, followed by three 2D convolutional/max-pooling layers, a flattening step, and finally a set of dense layers. After output generation (e.g., OFF), a gradient-weighted activation mapping is performed to visualize activation maximization regions responsible for delivered predictions (VIS4Map). (B) Histograms of the percentage overlap between VIS4Maps generated from a CNN pre-trained to predict minimum free energy (MFE) using 120-nt RNA sequences and MFE maps generated by NUPACK. When analyzed using 500 random test set sequences, the distributions of correctly matched and randomly assigned maps are distinct with increased percentage overlap from matched samples as compared to unmatched. (C) Examples of saliency VIS4Maps compared with their corresponding MFE structures as predicted by NUPACK for three randomly selected 60-nt RNA sequences. See Fig. S11A for additional examples with 120-nt RNA sequences. (D) Four representative VIS4Map examples of randomly selected 118-nt RNA toehold switch sequences from an OFF-predictive CNN model. (E) Averaged VIS4Maps of 10,125 randomly selected toehold switch RNA sequences from our library test-set processed with our OFF-predicting CNN model (left) and compared their corresponding averaged MFE maps obtained using NUPACK (right). (F) Averaged VIS4Maps of the 10% most accurately predicted switches sorted by quartile from lowest OFF (tight) to highest OFF (leaky), inset for the toehold and the hairpin stem. After contrast enhancement of averaged VIS4Maps to visualize sparsely distributed secondary structures, a noticeable increase in structures outside of the prominent equilibrium-designed switch hairpin structure appears to correlate with increased toehold leakiness. A toehold switch schematic (right) is shown to denote how incorrectly folded and potentially weaker kinetically stable intermediate structures might compete with the correctly folded structure that is designed to be reached at equilibrium.

To validate the feasibility of our visualization approach, we first pre-trained a CNN to predict NUPACK MFE values from complementarity map representations of a randomly selected *in silico* RNA sequence dataset. Because MFE is directly determined by RNA secondary structure, we anticipated that a CNN undergoing this pre-training would likely pay attention to secondary structure features, a situation that was confirmed through visualization of individual attention maps (Fig. 5B,C). Additionally, we found that the use of a complementarity map input improved the CNN’s predictions of MFE from R^2^=0.6 to R^2^=0.74 compared with a one-hot sequence input (Fig. S11). Indeed, the saliency maps generated from a CNN trained on a complementarity map input contained primarily diagonal features that showed a statistically significant degree of agreement with the MFE structures from which NUPACK based its MFE calculations (Fig. 5B,C, S11). Hence without prior knowledge of the algorithm or parameters NUPACK uses to calculate MFE, our CNN was able to learn similar abstractions as NUPACK, which we then used to intuitively visualize underlying relevant RNA secondary structures utilizing our complementarity map input representation. We named this approach for interpreting RNA deep learning models <u>Vis</u>ualizing <u>S</u>econdary <u>S</u>tructure <u>S</u>aliency <u>Map</u>s or VIS4Map.

Encouraged by our CNN’s ability to elucidate RNA secondary structure features directly from training data, we applied VIS4Map to our entire toehold switch dataset. When trained on a complementarity map representation (Fig. 5D) both in regression mode and classification mode, VIS4Map significantly outperformed an MLP trained on rational thermodynamic features; however, VIS4Map did not significantly outperform our MLP trained on one-hot inputs, similar to the case of our other higher capacity models (Fig. 4A,B,C,D). Encouragingly, nonetheless, we found that saliency maps produced by this CNN model displayed clear diagonal secondary structure features (Fig. 5D). These structures appear to span from hybridization between the toehold and the ascending stem, to hybridization between the descending stem and the downstream linker. We confirmed the biological relevance of these features by averaging saliency maps and finding that the shared structures corresponded to the designed on-target structure of the switch hairpin (Fig. 5E). We further analyzed learned features outside of the designed equilibrium structure by sorting saliency maps using the toehold switch OFF signal (Fig. 5F, S12). We found that for leakier (high OFF) switches, the CNN identified a high degree of salient off-target secondary structures that could compete with the main hairpin stem and thereby exposed the RBS, whereas for tight (low OFF) switches the CNN identified fewer competing off-target secondary structures. In the context of general riboregulator behavior, these findings support the hypothesis that leaky expression from an RBS repressed by secondary structures can be caused by the misfolding of the repressive structure into less stable kinetic intermediate conformations (*29, 32*) (Fig. 5F, right).

The fact that VIS4Map was able to identify both equilibrium and kinetically stable RNA secondary structures indicates a remarkable ability to uncover biologically relevant information, which in this case supports currently postulated hypotheses on prokaryotic translation initiation. Importantly, the identified secondary structure features could not have been visualized using the one-hot sequence representation commonly associated with saliency maps (*10, 11, 18, 20*). These findings compound to the advantage of using sequence-only deep learning approaches for analyzing RNA synthetic biology tools. Outside of toehold switches and other synthetic RNA systems, we anticipate VIS4Map will be broadly useful for the discovery of previously unknown equilibrium or kinetically stable structures contributing to RNA biology, that are not predicted by current mechanistic RNA structure models.

## Discussion

Here we presented a high-throughput DNA synthesis, sequencing, and deep learning pipeline for the design and analysis of a synthetic system in RNA biology. Having produced a toehold switch dataset ~100-fold larger than previously published as a model system for investigating synthetic RNA response elements (*2–6, 15, 16*), we demonstrated the benefits of using deep learning methods that directly analyze sequence rather than relying on calculations from mechanistic thermodynamic and kinetic models. This approach resulted in a tenfold improvement in functional prediction R^2^ over an ensemble of commonly used thermodynamic and kinetic features. Moreover, the validation of our deep learning models on an external previously characterized dataset, as well as the holdout prediction of every individual viral genome in our dataset, further demonstrated the robust biological generalization of our models.

As with most work in RNA synthetic biology, all previous attempts to improve toehold switch functionality have relied on the guidance of mechanistic thermodynamic modeling and low-throughput datasets (*2–8, 15, 16*). Too frequently, rational design rules fail to give meaningful predictions of function for RNA-based synthetic systems. The results presented here suggest that the biological processes underlying RNA biology may be more complex than current state-of-the-art analyses take into account and that high-throughput DNA synthesis, sequencing, and deep learning pipelines can be more effective for modeling said complexity. Combining improved predictions with enhanced understanding, our novel VIS4Map method further allowed us to visualize the equilibrium and kinetic secondary structure features that our deep learning models identified as important to the leakage of the switch OFF state. While secondary structures identified by NUPACK, Kinfold, and other rational mechanistic models are limited by predefined abstractions, which may cause significant information loss, our approach explored sequence space in an unrestricted manner and analyzed all possible RNA secondary structures. VIS4Map could prove useful for identifying complex secondary structure information that might otherwise be ignored by simplified physical energetic models of RNA folding.

The dataset reported here also represents an extensive repository of characterized toehold switches, which could be used to accelerate the development of future cell-free diagnostics (*3, 4, 15, 16*). These switches tile the entire genomes of 23 pathogenic viruses of high clinical importance, as well as tiling hundreds of human transcripts, including many that are differentially expressed in cancerous phenotypes (*42, 43*). The total cost of our flow-seq pipeline equates to ~$0.08 per measurement, suggesting that the benefits of high-throughput design and assaying of RNA synthetic biology tools could be made widely accessible. We hope that this work will encourage the use of high-throughput data collection for the training of deep learning systems, paired with more interpretable neural network architectures unrestricted by thermodynamic or kinetic secondary structure models for improved prediction and insight generation in RNA synthetic biology.

## Acknowledgments

We thank Diogo Camacho for his aid in statistical analysis and data preparation methodologies. We also thank Max A. English, Nina M. Donghia and Peter Q. Nguyen for their support in experimental activities of this work. We also thank Timothy Kassis for the helpful discussions and advice relating to the implementation of deep learning model architectures. We also thank Max Schubert and Pierce Ogden for advice in optimizing library cloning. We also thank Howard Salis for giving us access to his RBS Calculator 2.1 API.

## Author contributions

N.A.M and A.S.G designed, constructed, and tested the toehold switch database; N.A.M, A.S.G, and L.R.S., planned and performed experiments, wrote code, analyzed the data, and wrote the manuscript; J.J.C. and G.C. directed overall research and edited the manuscript.

## Data and materials availability

All data needed to evaluate the conclusions in the paper can be found in the paper and/or the Supplementary Materials. Correspondence and requests for materials should be addressed to J.J.C.

## Additional information

Supplementary Methods

Supplementary Text

Figures S1-S11

Tables S1-S4

## References

1. F. J. Isaacs, D. J. Dwyer, J. J. Collins, RNA synthetic biology. Nature biotechnology 24, 545 (2006).

2. A. A. Green, P. A. Silver, J. J. Collins, P. Yin, Toehold switches: de-novo-designed regulators of gene expression. Cell 159, 925–939 (2014).

3. K. Pardee et al., Rapid, low-cost detection of Zika virus using programmable biomolecular components. Cell 165, 1255–1266 (2016).

4. M. K. Takahashi et al., A low-cost paper-based synthetic biology platform for analyzing gut microbiota and host biomarkers. Nature communications 9, 3347 (2018).

5. A. A. Green et al., Complex cellular logic computation using ribocomputing devices. Nature 548, 117 (2017).

6. S.-J. Kim, M. Leong, M. B. Amrofell, Y. J. Lee, T. S. Moon, Modulating responses of toehold switches by an inhibitory hairpin. ACS synthetic biology 8, 601–605 (2019).

7. M. Krishnamurthy et al., Tunable riboregulator switches for post-transcriptional control of gene expression. ACS synthetic biology 4, 1326–1334 (2015).

8. J. Kim et al., De-Novo-Designed Translational Repressors for Multi-Input Cellular Logic. bioRxiv, 501783 (2018).

9. A. C.-Y. To et al., A comprehensive web tool for toehold switch design. Bioinformatics 34, 2862–2864 (2018).

10. H. K. Kim et al., Deep learning improves prediction of CRISPR–Cpf1 guide RNA activity. Nature biotechnology 36, 239 (2018).

11. S. Webb, Deep learning for biology. Nature 554, (2018).

12. C. Angermueller, T. Pärnamaa, L. Parts, O. Stegle, Deep learning for computational biology. Molecular systems biology 12, (2016).

13. M. Wainberg, D. Merico, A. Delong, B. J. Frey, Deep learning in biomedicine. Nature biotechnology 36, 829 (2018).

14. D. M. Camacho, K. M. Collins, R. K. Powers, J. C. Costello, J. J. Collins, Nextgeneration machine learning for biological networks. Cell 173, 1581–1592 (2018).

15. K. Pardee et al., Paper-based synthetic gene networks. Cell 159, 940–954 (2014).

16. D. Ma, L. Shen, K. Wu, C. W. Diehnelt, A. A. Green, Low-cost detection of norovirus using paper-based cell-free systems and synbody-based viral enrichment. Synthetic Biology 3, ysy018 (2018).

17. G. Chuai et al., DeepCRISPR: optimized CRISPR guide RNA design by deep learning. Genome biology 19, 80 (2018).

18. J. Luo, W. Chen, L. Xue, B. Tang, Prediction of activity and specificity of CRISPR-Cpf1 using convolutional deep learning neural networks. BMC bioinformatics 20, 332 (2019).

19. S. Zhang, H. Hu, T. Jiang, L. Zhang, J. Zeng, TITER: predicting translation initiation sites by deep learning. Bioinformatics 33, i234–i242 (2017).

20. J. Zuallaert, M. Kim, Y. Saeys, W. De Neve, in 2017 IEEE International Conference on Bioinformatics and Biomedicine (BIBM). (IEEE, 2017), pp. 1233–1237.

21. E. C. Alley, G. Khimulya, S. Biswas, M. AlQuraishi, G. M. Church, Unified rational protein engineering with sequence-only deep representation learning. bioRxiv, 589333 (2019).

22. D. B. Goodman, G. M. Church, S. Kosuri, Causes and effects of N-terminal codon bias in bacterial genes. Science 342, 475–479 (2013).

23. J. N. Zadeh, B. R. Wolfe, N. A. Pierce, Nucleic acid sequence design via efficient ensemble defect optimization. Journal of computational chemistry 32, 439–452 (2011).

24. R. M. Dirks, M. Lin, E. Winfree, N. A. Pierce, Paradigms for computational nucleic acid design. Nucleic acids research 32, 1392–1403 (2004).

25. R. Lorenz et al., ViennaRNA Package 2.0. Algorithms for molecular biology 6, 26 (2011).

26. H. M. Salis, E. A. Mirsky, C. A. Voigt, Automated design of synthetic ribosome binding sites to control protein expression. Nature biotechnology 27, 946 (2009).

27. A. Espah Borujeni et al., Precise quantification of translation inhibition by mRNA structures that overlap with the ribosomal footprint in N-terminal coding sequences. Nucleic acids research 45, 5437–5448 (2017).

28. A. Espah Borujeni, A. S. Channarasappa, H. M. Salis, Translation rate is controlled by coupled trade-offs between site accessibility, selective RNA unfolding and sliding at upstream standby sites. Nucleic acids research 42, 2646–2659 (2013).

29. A. Espah Borujeni, H. M. Salis, Translation initiation is controlled by RNA folding kinetics via a ribosome drafting mechanism. Journal of the American Chemical Society 138, 7016–7023 (2016).

30. B. Reeve, T. Hargest, C. Gilbert, T. Ellis, Predicting translation initiation rates for designing synthetic biology. Frontiers in bioengineering and biotechnology 2, 1 (2014).

31. M. M. Meyer, The role of mRNA structure in bacterial translational regulation. Wiley Interdisciplinary Reviews: RNA 8, e1370 (2017).

32. S. Badelt, S. Hammer, C. Flamm, I. L. Hofacker, in Methods in enzymology. (Elsevier, 2015), vol. 553, pp. 193–213.

33. B. Sauerwine, M. Widom, Kinetic Monte Carlo method applied to nucleic acid hairpin folding. Physical Review E 84, 061912 (2011).

34. V. I. Jurtz et al., An introduction to deep learning on biological sequence data: examples and solutions. Bioinformatics 33, 3685–3690 (2017).

35. X.-Q. Liu, B.-X. Li, G.-R. Zeng, Q.-Y. Liu, D.-M. Ai, Prediction of Long Non-Coding RNAs Based on Deep Learning. Genes 10, 273 (2019).

36. J. Baek, B. Lee, S. Kwon, S. Yoon, Lncrnanet: long non-coding rna identification using deep learning. Bioinformatics 34, 3889–3897 (2018).

37. G. Aoki, Y. Sakakibara, Convolutional neural networks for classification of alignments of non-coding RNA sequences. Bioinformatics 34, i237–i244 (2018).

38. A. Fiannaca, M. La Rosa, L. La Paglia, R. Rizzo, A. Urso, nRC: non-coding RNA Classifier based on structural features. BioData mining 10, 27 (2017).

39. N. Frosst, G. Hinton, Distilling a neural network into a soft decision tree. arXiv preprint arXiv:1711.09784, (2017).

40. P. K. Koo, S. R. Eddy, Representation Learning of Genomic Sequence Motifs with Convolutional Neural Networks. BioRxiv, 362756 (2018).

41. K. Simonyan, A. Vedaldi, A. Zisserman, Deep inside convolutional networks: Visualising image classification models and saliency maps. arXiv preprint arXiv:1312.6034, (2013).

42. A. Dhawan, J. G. Scott, A. L. Harris, F. M. Buffa, Pan-cancer characterisation of microRNA across cancer hallmarks reveals microRNA-mediated downregulation of tumour suppressors. Nature communications 9, 5228 (2018).

43. Y. Xin-wei et al., STAT3 overexpression promotes metastasis in intrahepatic cholangiocarcinoma and correlates negatively with surgical outcome. Oncotarget 8, 7710 (2017).

