## Supplementary Information for "Deep Learning for RNA Synthetic Biology"

**This PDF file includes:**

Materials and Methods
Supplementary Figures (S1 to S13)
Supplementary Tables (S1 to S4)

### MATERIALS AND METHODS

#### Toehold Switch Architecture Selection

The first-generation toehold switch architecture from Green et al. (*1*) was selected in order to maximize the sequence variability in switch regions contributing to secondary structure. Where in later designs the trigger RNA only unwound a fraction of the stem (*1-3*), in this earlier design the entire hairpin stem was variably complementary to the trigger, increasing the diversity of characterized RNA hairpins (Fig. 1A). An alternative fused ON state was also utilized. Normally, toehold switches detect the presence of a separate trigger RNA transcribed *in trans* to the OFF-state switch mRNA. However, for the testing of a large library of toehold-switch pairings, a two-plasmid system becomes intractable because each switch is designed around a specific cognate trigger. A two-plasmid system can also increase stochasticity caused by copy number variability. Green et al. (*1*) found a strong positive correlation between conditions when the trigger is fused to the switch and conditions when un-fused, separate triggers are transcribed in excess. We confirmed this correlation ourselves on a subset of twenty toehold switches by comparing the signal from the alternative fused ON state used in our library to the measured ON/OFF from Green et al. (*1*). Green et al. did not report separate ON and OFF measurements, but have stated that due to a low switch plasmid copy number their OFF state rarely exceeded background autofluoresence, meaning that their reported ON/OFF ratios are approximations of ON state measurements. The resulting comparison of signal from the alternative fused ON state we measured and the un-fused ON/OFF ratio measured by Green et al using a two-plasmid system resulted in a Pearson R=0.8567, as seen in Fig. S1B. Thus, the ON state of the switch can be reliably approximated by fusing the trigger RNA to the 5' end of the switch mRNA using a constant, unstructured linker sequence (Fig. 1A, S1A), allowing for the direct synthesis of trigger-switch cognates on a single plasmid.

#### Library Trigger Sequence Selection

Viral genomes were obtained on November 6th, 2018, from https://www.ncbi.nlm.nih.gov/genome/viruses/ Each retrieved genome was tiled 30bp at a time (the trigger length), with a stride of 5bp, spanning the respective genome. Human transcription factors were obtained using ENSEMBL 94 BioMart (*4*) utilizing the Gene Ontology term GO:0044212 (transcription regulatory region DNA binding). The coding region of each transcription factor was tiled 30bp at a time with a stride of 10. A remaining portion of the designs (~10,000) was based on random 30bp triggers.

#### Toehold Library Synthesis

We designed 244,000 toehold switch variants using 230bp oligos, which were ordered and synthesized by Agilent. For each toehold switch variant, the oligo was designed containing the following sequence components in order from 5' to 3': 20nt of common backbone, a T7 Promoter, the 30nt Trigger sequence, a 20nt unstructured Linker, the 12nt Toehold, the 18nt Ascending Stem, a 11nt SD-containing Loop, the 18nt Descending Stem including the start codon, a 21nt AA-Linker, and the first 15nt of the GFP gene. A schematic of the design can be found in Fig. S1A. In the previous validation of the fused trigger approach by Green et al. (*1*), only part of the trigger was fused to avoid recombination of long repeated sequences, but the nature of our flow-seq pipeline allowed us to avoid this issue since the integrity of all variants was confirmed after measuring fluorescence through next-generation sequencing (NGS). The oligos were received at a stock amount of 10pmol, which we diluted in 500uL TE buffer for a working concentration of 20nM. Of this working stock, 0.25uL was used in 50uL qPCR reactions using NEB Q5 polymerase 2xMM with 50nM final concentration of appropriate primers. Two separate amplifications were done from the working stock of the oligo library for the ON and OFF states, respectively. One amplification, for the ON state, used a primer hybridizing to the 5' common backbone region. The resulting insert contained both the Switch RNA module and the Trigger attached to its 5' end. The second amplification, for the OFF state, used a primer hybridizing to the 20nt unstructured Linker and included a T7 promoter and the 5' common backbone region in its tail. The OFF-state insert contained only the Switch RNA module without the Trigger module attached. See Fig. S1A for a full schematic of the amplification scheme. A third amplification linearized a ColE1 plasmid backbone for subsequent ligation. This backbone was the same ColE1 backbone as was used in Green et al. (*1*) for transcribing trigger RNAs, but with a GFPmut3b-ASV gene inserted. All amplicons were cleaned from their reaction buffers by using carboxyl-coated magnetic beads (*5*) (protocol 4.3): 1x concentration of beads to clean the longer linear backbone product, and 2x bead concentration to clean the smaller insert products. Both inserts were ligated separately into the ColE1 backbone in front of the GFPmut3b-ASV gene using golden gate cloning, as follows. The linearized plasmid backbone was diluted to 500ng total mass. The ON or OFF insert was added according to a 1:1 molar ratio of insert to plasmid backbone. The inserts and backbone dilutions were prepared into 50uL ligation reaction volumes, containing 5uL NEB buffer 3.1, 5uL T4 ligase buffer, 1uL BsmBI, 0.5uL Dpn1, 1uL T4 ligase, and any remaining volume with nuclease-free water. The 50uL reaction was placed into a thermocycler for 100 cycles of two steps: 16C for 10 min and 37C for 10 min. A final enzyme inactivation step at 65C for 15min was done. The ligation products were precipitated out of their reaction buffers using ethanol precipitation. The 50uL ligation reactions were added to 1.5mL Eppendorf tubes containing 150uL of pure ethanol, 5uL 0.3M sodium acetate (pH 5.2), and 1uL glycoblue. Tubes were left on dry ice for 20 min and then immediately placed in a 4C tabletop centrifuge and spun at max RPM for 30 min. Tubes were decanted, and 175uL of 70% ethanol was added to the tube containing the pellet. Tubes were spun at max speed for 5 min. Tubes were then removed from the centrifuge, decanted, and allowed to dry for 15 min. Ligation products were then eluted in 4uL TE buffer. For initial library transformation, 50uL EclonI Supreme cells were given the full 4uL ligation product elution and electro-transformed. Transformation efficiencies exceeding $10^7$ CFU/mL were achieved, and the expanded cells were harvested using a MaxiPrep kit (Qiagen). The resulting pool of plasmids was then electroporated into BL21 star *E. coli*, where transformation efficiencies exceeding $10^6$ were achieved.

**Flow-seq Pipeline**

Induction was achieved by expanding BL21 cells overnight at 37C in LB media with carbenicillin (carb) selection and then diluted 50x into fresh media. After the cells reached OD600 of 0.3 at 37C (~2 hours of growth), 0.2mM IPTG was added, and the cells were allowed to express for another 3 hours at 37C. The cells were then moved to room temperature and sorted on a Sony SH800 FACS machine with four bins. A positive control consisting of Switch #4 from Green et al. (*1*), one of the highest performing switches from that study's first-generation design, was cloned both in its OFF state and in the modified fused-trigger ON state. This positive control switch was then used to mark the highest and middle bins of GFP signal, while a negative control consisting of a pUC19 plasmid (containing no GFP) was used to mark the lowest bin of GFP signal (Fig. S2). Approximately 40 million events were sorted for each library. Cells in collected bins were diluted 10x into fresh LB media with carb selection and allowed to expand overnight at 30C. The expanded cells were then harvested using a MaxiPrep kit (Qiagen).

**Deep Sequencing, Read Data Processing and Read Count Analysis**

Plasmid collected from sorted cells was amplified using NEB Q5 polymerase 2xMM and primers targeting the common backbone region upstream and downstream of the variable toehold region. The resulting 184bp (OFF) or 224bp (ON) PCR products were then analyzed by NGS using a MiSeq or NextSeq instrument (Illumina). Raw paired-end sequencing reads were quality filtered and merged with PEAR 0.9.1. Only sequences matching our intended designs were retained for further analysis. For the ON and OFF libraries, respectively, 10,390,207 reads and 20,788,966 reads were mapped to a correct switch sequence. The individual fluorescence distribution of the ON and OFF state for each switch was measured by calculating its frequency in each bin and assigning a normalized signal metric in the range of [0,1] (Fig. 1C, D), and an ON/OFF metric was calculated as the difference between the ON and OFF signal metrics independently (Fig. 1E). Frequencies of each variant were tabulated for each cell-sorted bin and normalized to the total reads per bin. Each variant's functional value was computed as the weighted mean of its normalized frequencies across all bins, scaled between 0 and 1. The ON/OFF ratios were then calculated as the subtracted difference between ON and OFF (since the fluorescence data had been collected on a logarithmic scale), resulting in a range scaled between -1 and 1.

**Library Quality Control**

A second biological replicate of our flow-seq pipeline was carried out that produced 60,800 ON measurements, 98,295 OFF measurements, and 30,101 ON/OFF ratio measurements where both ON and OFF were available for the same switch. The $R^2$ and MAE between our two datasets were calculated at different read count thresholds. Based on the results (Fig. S3), five different QC thresholds were established, some of which also included standard deviation cutoffs (Table S1, Fig. S13). QC1 and QC2 contained OFF data with significantly worse $R^2$ compared to QC3, QC4, and QC5, but only QC1 contained OFF data with worse MAE. We determined that the inter-replicate drop in $R^2$ for OFF values was mainly due to the skewness of the data – indeed, the OFF data consistently showed worse $R^2$ values than the ON data throughout the paper, despite having consistently better MAE values. Therefore, we chose to trust in the inter-replicate MAE values more than the inter-replicate $R^2$ metric for the OFF data.

To further evaluate the different QC levels, the most stringent data (QC5) were withheld as a test set, and an MLP fed a one-hot representation of the toehold sequence was trained on the four lower QC levels. The results for both predictive $R^2$ and MAE showed QC1 to be of significantly inferior quality, but QC2, QC3, and QC4 to be of roughly similar quality (Fig. S4). This result was consistent with the fact that inter-replicate MAE was notably worse at the QC1 count threshold but essentially unchanged across the read count thresholds contained by QC2, QC3, and QC4. The QC2 dataset gave the best predictive results by a small margin and was also significantly larger than QC3 or QC4 (Table S1). With these analyses in mind, QC2 was chosen as the final threshold for inclusion in our dataset. Within the measured ON/OFF ratios in the QC2 dataset, 40,824 had triggers of viral origin, 47,005 had triggers of human origin, and 3,705 had randomly generated trigger sequences.

### Cell-Free Switch Validation

Eight of the best performing switches (ON/OFF>0.97) and eight of the worst performing switches (ON/OFF<0.05) were synthesized as PCR products, as previously described (*2*). Briefly, they were ordered as single Ultramer oligos (IDT) without the Trigger fused, from the T7 promoter to the first 36nt of the common linker and GFP sequences. These were added to a GFP gene by a single PCR amplification step. Triggers were *in vitro* transcribed from separate oligos that contained the antisense sequence and the antisense T7 promoter, to which the sense strand of the T7 promoter was annealed. Trigger RNA was purified using an RNA Clean & Concentrator kit (Zymo), while Switch DNA was purified using a MinElute kit (Qiagen). To a 5uL PURExpress reaction were added 2U/uL Murine RNAse Inh, 5nM of Toehold Switch PCR product, and either no Trigger RNA or 10uM of Trigger RNA. Measurements of GFP velocity can be found in Fig. S5. The exact Switches tested and their library assay measurements can be found in Table S2.

### Calculations Made with ViennaRNA, Kinfold, and the RBS Calculator

All thermodynamic MFE and ensemble defect calculations, as well as kinetic Kinfold calculations, were obtained using a custom-made python code including libraries from packages such as Biopython (Ref: https://github.com/biopython/biopython), ViennaRNA (Ref: https://github.com/ViennaRNA/ViennaRNA), RNAsketch (Ref: https://github.com/ViennaRNA/RNAsketch) and Pysster (Ref: https://github.com/budach/pysster). Calculations of thermodynamic rational parameters to include in our database were obtained from toehold RNA sequences by taking each basal 145-nucleotide toehold sequence and then isolating different sections (e.g., GGG, Trigger, Loop1, Switch, Loop2, Stem1, AUG, Stem2, Linker, Post-linker) into distinct sub-sequences with biological relevance for functional analysis (see Fig. S1, Table S4). Minimum Free Energy (MFE) was calculated for all these sections using the previously reported python-based ViennaRNA Library. MFE calculation using ViennaRNA also specifies a secondary structure in dot-parens-plus notation (unpaired base = dot, base-pair = matching parentheses, and nick between strands = plus). Ideal structures are assumed to be connected and free of pseudoknots. These ideal secondary structures for such sections are:

```
SwitchOFF = '..............................(((((((((...((((((...........))))))...)))))))))'
SwitchOFF_GFP = '..............................(((((((((...((((((...........))))))...)))))))))..(((.......(((((.....)))))..)))....'
SwitchOFF_NoTo = '(((((((((...((((((...........))))))...)))))))))..(((.......(((((.....)))))..)))....'
SwitchON = '...((((((((((((((((((((((((((((((.....................))))))))))))))))))))))))))))))............................'
SwitchON_GFP = '...((((((((((((((((((((((((((((((.....................))))))))))))))))))))))))))))))..............................(((.......(((((.....)))))..)))....'
ToeholdON = '.....................((((((((((((.....................))))))))))))'
Stem = '(((((((((...((((((...........))))))...)))))))))'
StemTop = '((((((...........))))))'
```

Ensemble defect as a rational parameter was calculated via ViennaRNA/NUPACK for each of the toehold switches in the above subsets of sequence regions: SwitchOFF, SwitchOFF_GFP, Switch_OFF_NoTo, SwitchON, SwitchON_GFP, ToeholdON, Stem, StemTop. This calculation used both the native (calculated from MFE) and the ideal (pre-defined above) dot-Bracket representation for each sequence to assess the average number of nucleotides that are incorrectly paired at equilibrium. Thirty rational parameters were calculated for each toehold using these methods (fourteen MFE values, eight ideal ensemble defect values, and eight native ensemble defect values).

Kinetic analyses using Kinfold were run from the ViennaRNA package. The OFF-switch sequence was selected, spanning nucleotides 50 to 134 in Table S4 from the start of the toehold to the end of the linker. Due to the large size of the toehold switch RBS, Kinfold trajectories ran for 100-1000x longer than for RBS's previously analyzed relating to the RBS calculator in Borujeni et al. (*6*) (Fig. S7b). Hence our analysis was scaled down to the QC4 dataset (containing 19,983 total switches), with 100 Kinfold trajectories run for each switch with a maximum stopping time of $10^3$ arbitrary Kinfold units (au). The energy and time at each step of each trajectory were recorded. If the MFE structure was reached within $10^3$ au, it was assumed that the RNA would remain in the MFE structure for the rest of the $10^3$ au timeframe. From each energy trajectory spanning $10^3$ au, the average energy (in kcal/mol) was calculated by integrating the energy-time curve and dividing by $10^3$. For each switch, the following features were extracted: the mean and standard deviation of the average energy of its 100 sampled trajectories (Fig. S7C), the ratio of the mean average energy to the MFE (Fig. S7E), and the fraction of trajectories that reached the MFE structure within the analyzed $10^3$ timeframe (Fig. S7D).

For predictions by the RBS Calculator, an API was used to access the most recent publicly available version (2.1). Due to limiting computational costs, the QC3 dataset was used instead of the QC2 dataset. For each switch, the translation initiation rate (TIR) of the on-target start codon was predicted for both the ON and OFF states ("SwitchON_GFP" and "SwitchOFF_GFP" respectively in Table S4).

**K-mer Motif Search**

In order to compare sequence-level motifs between the best and worst variants measured in our dataset, we performed a k-mer search for over-represented sequence motifs at the tails of our observed functional values. We first filtered the variants for high quality, retaining those with a QC4 score or above. We then took the top and bottom 1,000 variants based on the ON and OFF functional values, respectively. We utilized DREME (*7*) to test for enrichment or depletion of all possible subsequences of length 3-16 bases, using the indicated foreground and background frequencies. All results above the default E-value cutoff are shown (Fig. 3A, Table S3)

**Deep Learning Model Architectures**

*MLP – Rational Features*

The multilayer perceptron (MLP) model based on rational features included a 30-feature input followed by three dense fully connected layers of 25, 10, and 7 neurons, respectively, with rectified linear unit (ReLU) activation, batch normalization, and 20% dropout. This network was then fed to a final three-neuron layer (ON, OFF, ON/OFF) with linear activation for regression output, or to a final two-neuron layer (ON/OFF: binarized at +/- 0.7) with softmax activation for classification output.

*MLP – OneHot Seq*

The MLP model based on the one-hot encoded full 145-nucleotide sequence input was achieved by using a flatten layer followed by three dense layers with ReLU activation, batch normalization, and 20% dropout. Dense layers used 128, 64, and 32 neurons, respectively. This network was then fed to a final three-neuron layer (ON, OFF, ON/OFF) with linear activation for regression output, or to a final two-neuron layer (ON/OFF: binarized at +/- 0.7) with softmax activation for classification output.

*MLP – Hybrid Rational Features/OneHot Seq*
The ensemble MLP model was based on the rational features, as well as a one-hot encoded full 145-nucleotide sequence as input. To construct this model, two networks were assembled in parallel. The first network uses the same architecture for the MLP model with rational features, while the second network used the architecture of the MLP model for one-hot encoded 145-nucleotide sequences. Both networks were then concatenated and connected to a four-neuron dense fully connected layers with ReLU activation. This network was then fed to a final three-neuron layer (ON, OFF, ON/OFF) with linear activation for regression output, or to a final two-neuron layer (ON/OFF: binarized at +/- 0.7) with softmax activation for classification output.

*CNN – OneHot Seq*
The Convolutional Neural Network (CNN) model based on the one-hot encoded full 145-nucleotide sequence as input was achieved by direct feeding of the input to three convolutional layers with ReLU activation, batch normalization, and 20% dropout. The convolutional layers used had 32, 64, and 128 filters of size 3, respectively. Same-padding was used with L1 and L2 kernel regularization. The output from the convolutional layers was flattened and fed to two fully connected sequential dense layers of 16 neurons each with ReLU activation, batch normalization, and 20% dropout. This network was then fed to a final three-neuron layer (ON, OFF, ON/OFF) with linear activation for regression output, or to a final two-neuron layer (ON/OFF: binarized at +/- 0.7) with softmax activation for classification output.

*CNN – 2D Complementarity Map*
The Convolutional Neural Network (CNN) model based on the one-hot encoded categorical 2D complementarity-directional matrix from the full 145-nucleotide sequence as input was achieved by direct feeding of the input to three convolutional layers with ReLU activation, batch normalization, and 30% dropout. The convolutional layers used had 32, 64, and 128 filters of size 5x5 respectively. Same-padding was used with L1 and L2 kernel regularization. The output from the convolutional layers was flattened and fed to two fully connected sequential dense layers of 16 neurons each with ReLU activation, batch normalization, and 20% dropout. This network was then fed to a final three-neuron layer (ON, OFF, ON/OFF) with linear activation for regression output, or to a final two-neuron layer (ON/OFF: binarized at +/- 0.7) with softmax activation for classification output.

*LSTM – OneHot Seq*
The Long Short-Term Memory (LSTM) recurrent neural network model on the one-hot encoded full 145-nucleotide sequence as input was achieved by direct feeding of the input to a network with 128 recurrent units. The output of this was then connected to 100-neuron fully connected dense layer with ReLU activation, followed by batch normalization and 30% dropout. This network was then fed to a final three-neuron layer (ON, OFF, ON/OFF) with linear activation for regression output, or to a final two-neuron layer (ON/OFF: binarized at +/- 0.7) with softmax activation for classification output.

All models were trained using a maximum of 300 epochs, considering a 20-epoch early stopping patience, which gets triggered upon lack of model improvement on the validation set. Batch size for all models was 64*(1+ngpus), where ngpus is defined as the number of used graphic processing units during model training. All trained regression models were verified for reported metrics using 10-fold cross-validation, while classification-trained models were evaluated on three shuffled test sets as indicated.

**Complementarity Matrix and VIS4Map**

Complementary maps were defined as a One-Hot Encoded Categorical 2D Complementarity-directional Matrix (total number of tensor dimensions = 3) constructed by defining columns and rows of the matrix as the position of potential complementarity between any two given pairs of nucleotides in a single RNA sequence. The value in each position is defined as a one-hot encoded categorical variable according to the Watson-Crick pairing of the two nucleotides defining that position. Nucleotide pairings are assigned the following category: G-C (6) = [0 0 0 0 0 0 1], C-G (5) = [0 0 0 0 0 1 0], A-U (4) = [0 0 0 0 1 0 0], U-A (3) = [0 0 0 1 0 0 0], G-U (2) = [0 0 1 0 0 0 0], U-G (1) = [0 1 0 0 0 0 0], NonWCpairs (0) = [1 0 0 0 0 0 0]. VIS4Maps were generated using a modified algorithm, attention, activation maximization and saliency map visualization for Keras (Keras-Vis, Ref: `https://github.com/raghakot/keras-vis`) with tensorflow backend.

In this case, gradients were calculated from a regression model for all regions of the image to visualize what spatial features cause the predicted output to increase. To visualize the toehold regions that are mostly responsible for each prediction, small positive or negative gradients are highlighted using a normalization strategy. Given this information, such techniques allow us to generate heatmap-encoded saliency map images that spatially relate to the toehold regions in the complementarity map that lead to accurate predictions.

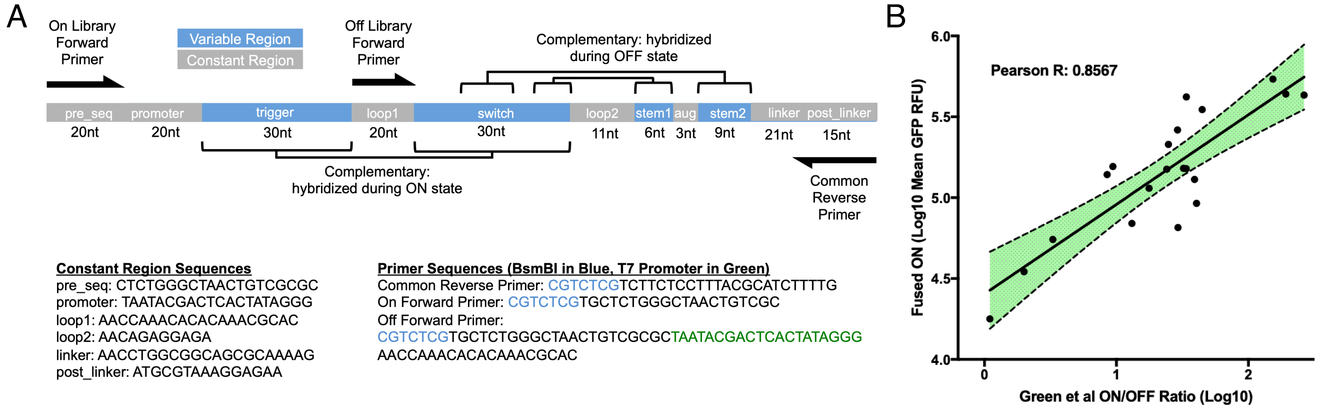

**Fig. S1. Design and validation of oligomer library.** Individual toehold switch constructs within the library were synthesized from a pool of oligomers, and a representative panel of constructs was verified against a previously published dataset. (A) Schematic of the pooled library oligo used for the synthesis of our high-throughput toehold switch library. Distinct toehold construct regions include: *pre_seq* (plasmid backbone sequence), *promoter* (T7 promoter including GGG), *trigger* (toehold-unique), *switch* (complete toehold and ascending stem), *loop1* (region linking *trigger* to *switch*), *loop2* (main toehold switch hairpin loop containing the RBS), *stem1* (top half of descending stem), *atg* (start codon), *stem2* (bottom half of descending stem), *linker* (21nt sequence of unstructured amino acids) and *post_linker* (first 15nt of GFP). Further detail can be found in Table S4. Amplification primers for both ON and OFF libraries (including the common reverse primer) are shown with black arrows. (B) Comparison of ON state GFP expression from a panel of 20 individually assayed switches from our high-throughput toehold switch pipeline against the ON/OFF ratio for equivalent switches reported by Green et al. (*1*). The agreement between the 5' fused triggers used in this work and the separately transcribed triggers used by Green et al. (*1*) was assessed based on the Pearson correlation coefficient (0.8567). GFP= Green fluorescent protein, nt=nucleotide, RBS=Ribosome binding site.

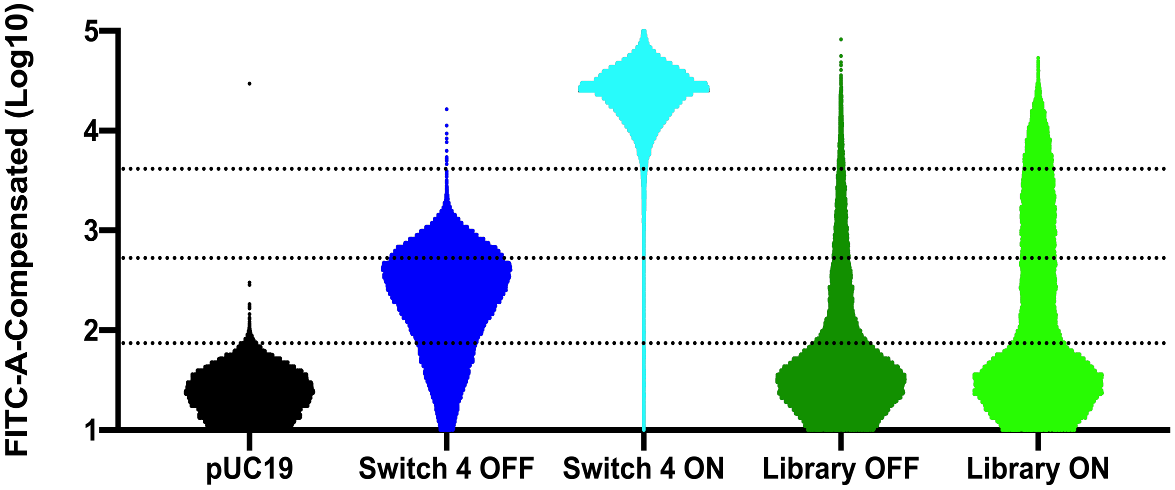

**Fig. S2. Library FACS distributions and empirically-derived sorting gates.** To determine the boundaries of the sorting gates for our high-throughput toehold switch pipeline, we used Switch #4 from Green et al. (*1*) in ON and OFF conformations as positive controls, and a pUC19 plasmid lacking a GFP gene as a negative control. Fluorescence distribution plots of IPTG-induced *E. coli* BL21-star cells from the three control conditions are shown alongside complete ON and OFF libraries for comparison. Boundaries for the four sorting bins are shown as dotted lines.

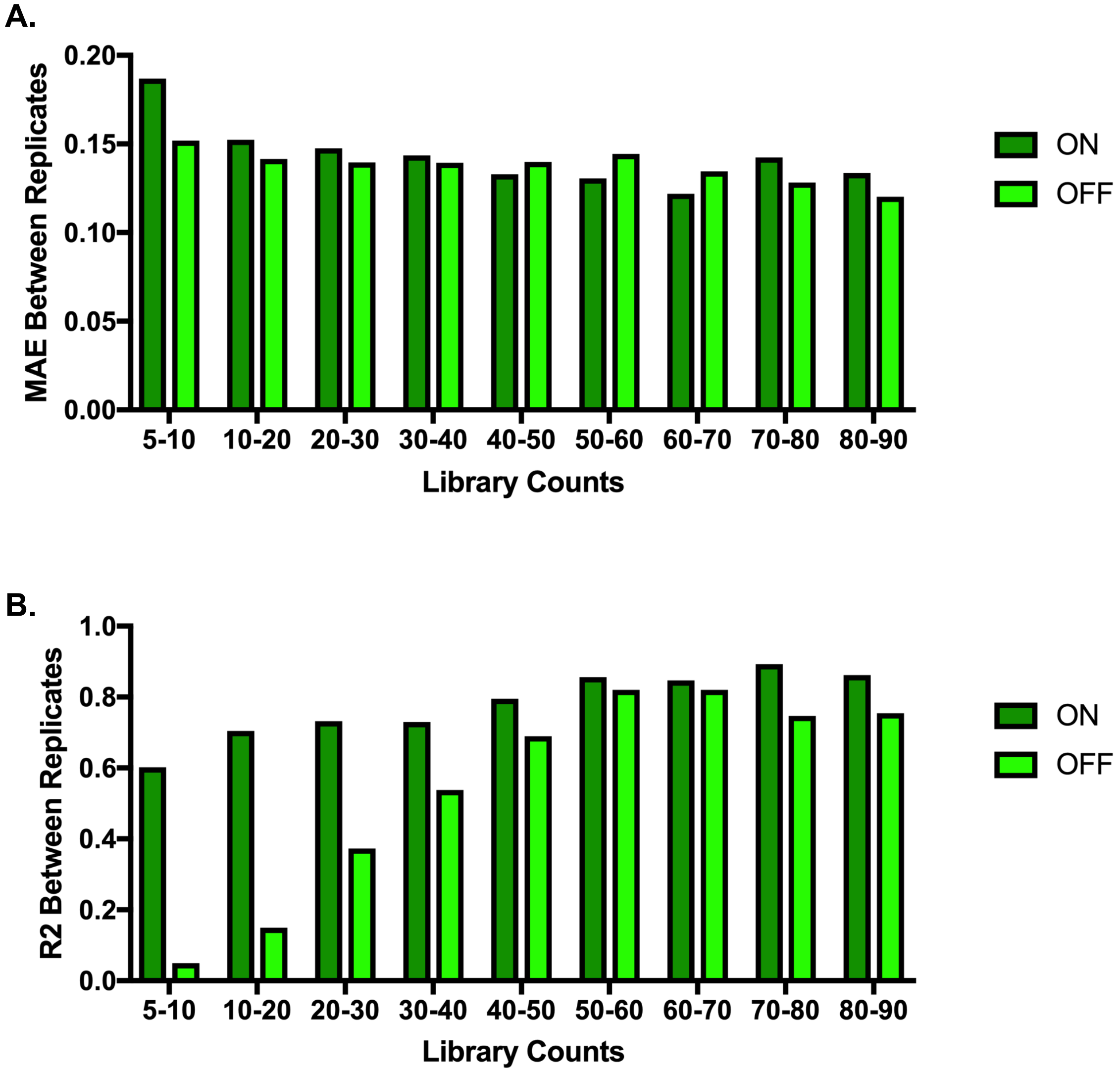

**Fig. S3. Inter-replicate variability of toehold switch libraries.** For the same initial toehold library, we performed two replicates of the BL21 transformation process followed by independent induction, sorting, and sequencing. Two metrics were used to compare the inter-replicate variability: (A) the mean absolute error (MAE), and (B) the $R^2$ correlation coefficient. Shown are the MAE and $R^2$ values for ON and OFF measurements at different ranges of library count thresholds.

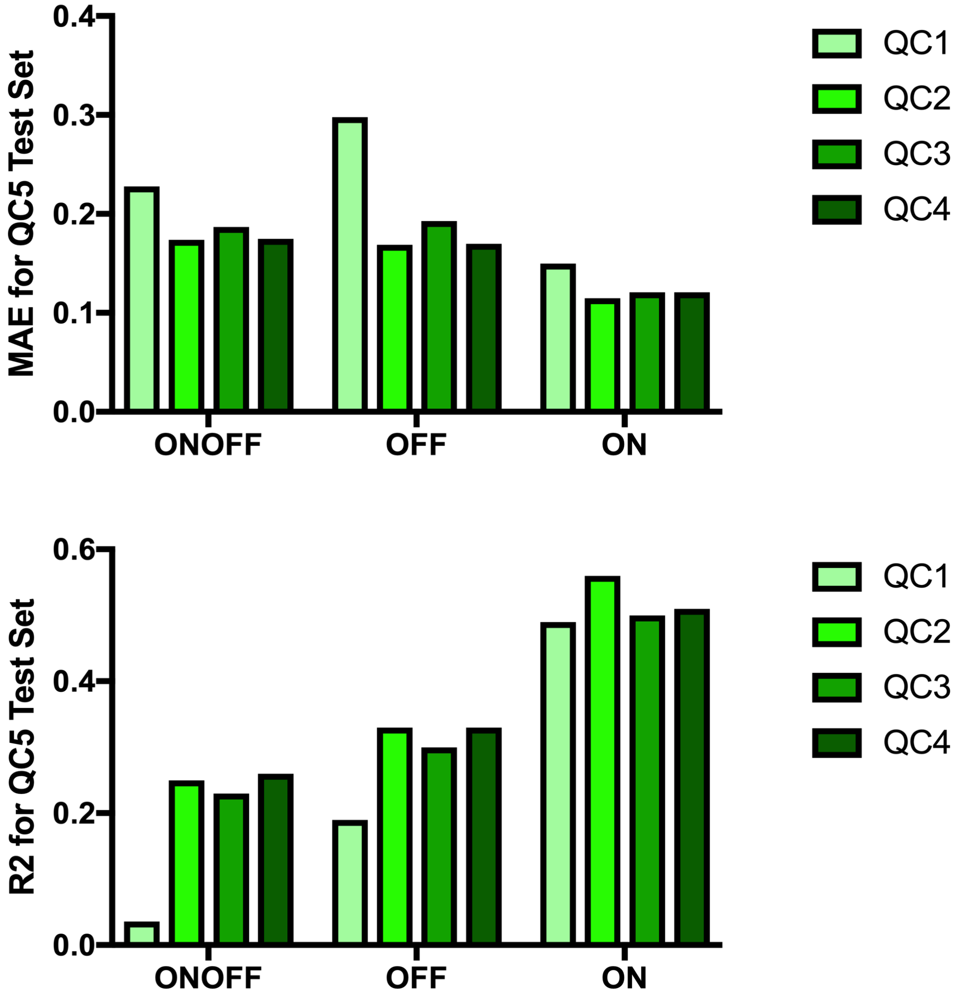

**Fig. S4. Effect of QC level on MLP performance.** The predictive power of our multilayer perceptron model was evaluated after training with datasets obtained from increasingly stringent quality control (QC) thresholds. The most stringent quality control group (QC5) was withheld as a test set, and an MLP trained on a one-hot representation of the toehold sequence was given either QC1, QC2, QC3, or QC4 as training data. From the resulting test-prediction of QC5 values, we show the MAE (upper panel), and the analogous $R^2$ correlation metric (lower panel) between the predicted and experimental values. See Table S1 for conditions for each QC level.

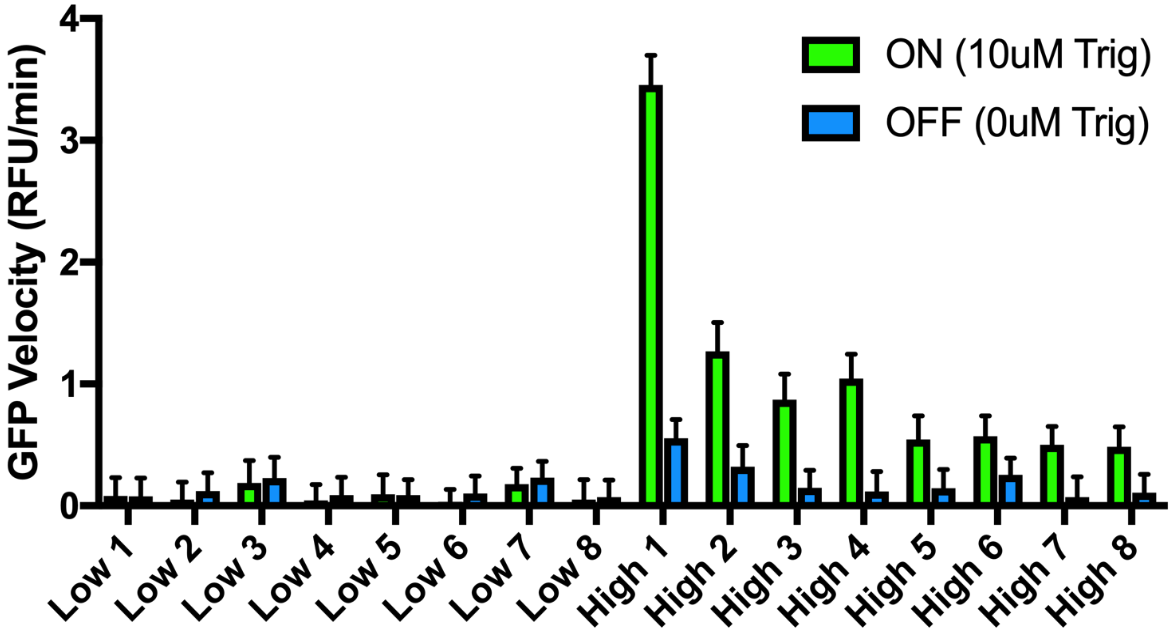

**Fig. S5. Cell-free toehold switch validation.** A panel of toeholds that showed either a low or high ON/OFF ratio as measured by our high-throughput flow-seq assay were individually cloned and assayed in a cell-free protein synthesis (CFPS) system. The timecourse velocities of GFP signal evolution are shown for the PURExpress CFPS reactions containing the sixteen switches with or without their separately transcribed RNA triggers. The sequences and flow-seq assay results for these sixteen switches can be found in Table S2.

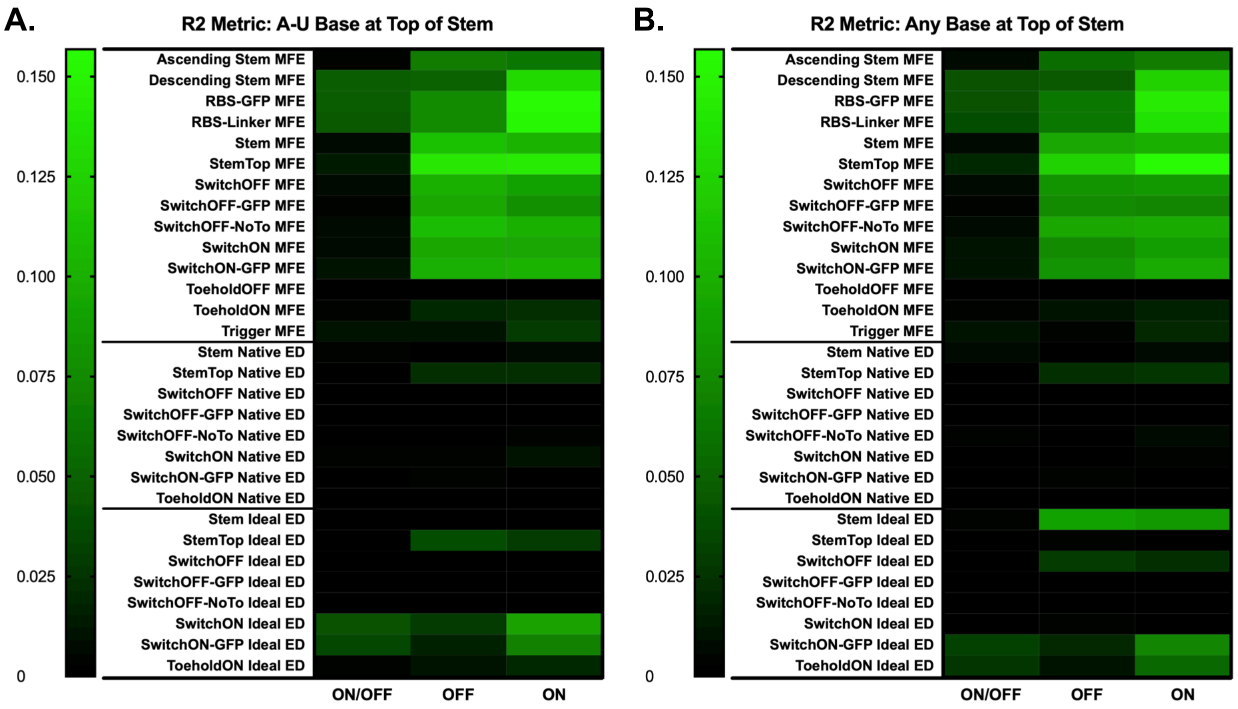

**Fig. S6. Correlation between rational thermodynamic features and toehold switch dataset, subsetted for A-U content.** We analyzed the $R^2$ coefficients between 30 commonly used thermodynamic features and the ON, OFF, or ON/OFF measurements of variants in our high-throughput dataset. (A) $R^2$ coefficients for the subset of switches that contained only an A-U or U-A base pair at the top of the toehold switch stem (positions 79 and 91 in Table S4). (B) $R^2$ coefficients for the entire set of switches, allowing for any base pair at the top of the toehold switch stem. Both $R^2$ value sets were compared to evaluate findings from Green et al. (*1*) where subsetting for switches with an A-U or U-A basepair at the top of the stem was sufficient to dramatically increase the predictive $R^2$ coefficient between thermodynamic features and measured ON/OFF. We found measurable differences between various thermodynamic features when subsetting for an A-U basepair at the top of the hairpin stem, particularly for those in the Ideal Ensemble Defect (ED) block. However, differences between the $R^2$ values in said subset and those obtained for other possible base-pairs were not statistically significant suggesting no overall increase in predictive value (p>0.05 for ON, OFF, and ON/OFF, two-tailed t-test).

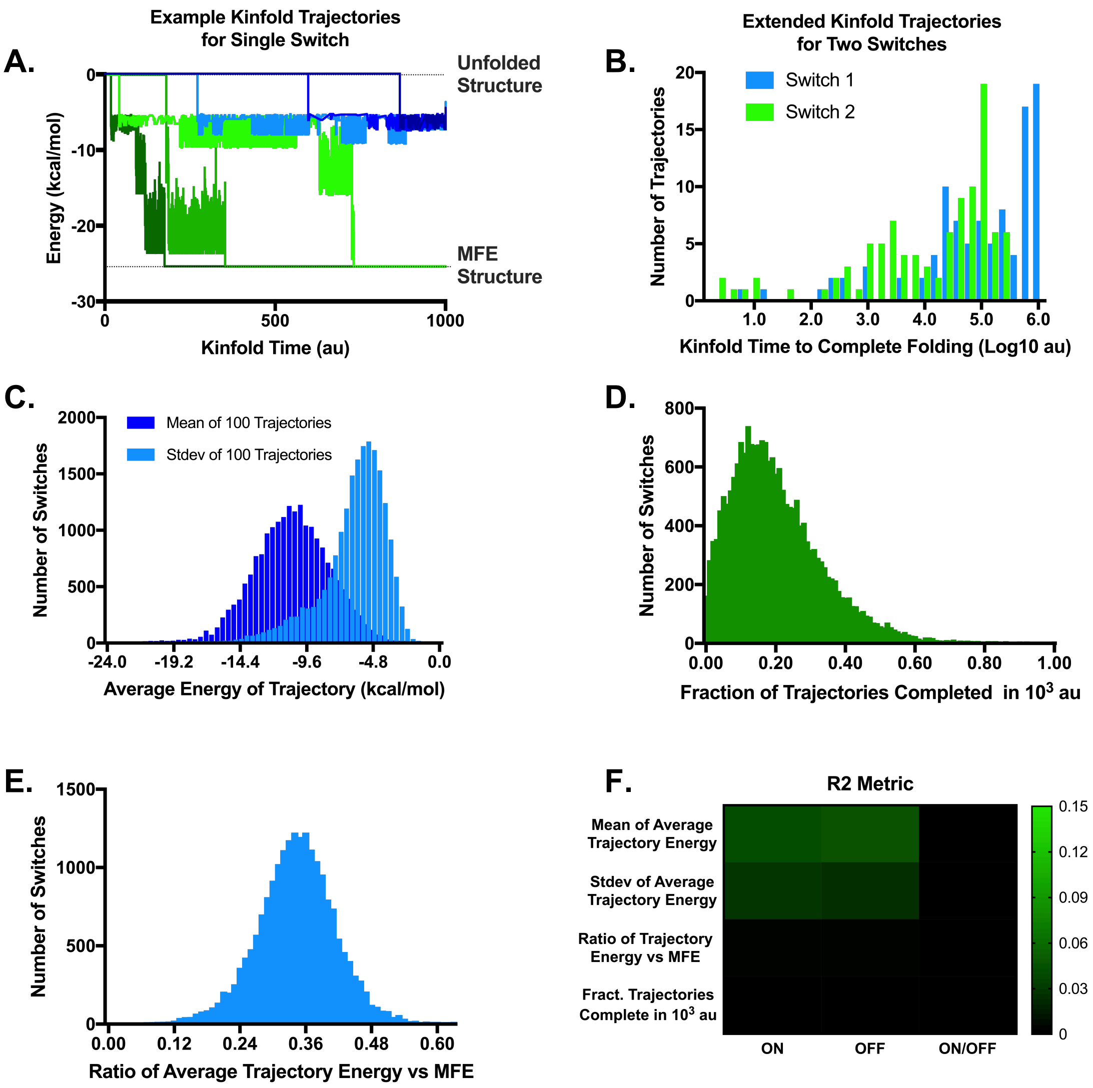

**Fig. S7. Kinetic toehold switch folding analysis using Kinfold.** Folding trajectories were run using the Kinfold package for the OFF-state switch sequence (positions 50-134nt in Table S4). (A) For a single representative toehold switch, six example trajectories are shown. Trajectories in green reached the MFE structure within $10^3$ arbitrary time units (au), while those in blue did not. (B) For two representative toehold switches, 100 trajectories were run for a maximum time of $10^6$ au. Histograms of the time required for a trajectory to reach the MFE structure are shown. Most trajectories took longer than $10^3$ au, compared to the Kinfold analyses in Borujeni et al. (*6*), where average trajectory times fell in the range of $10^1$-$10^3$ au, and $10^4$ au was the longest allowed trajectory time. (C,D,E,F) For each switch in the QC4 dataset (total 19,983 variants), 100 trajectories were run and the following measurements plotted: (C) histograms of the mean and negative standard deviation of the trajectories' average energy during the first $10^3$ au, (D) the fraction of trajectories that completed folding of the MFE structure before $10^3$ au, (E) the ratio of average trajectory energy to the minimum possible MFE energy, and (F) the $R^2$ correlation between the metrics in C,D,E and the empirical measurements in our toehold switch dataset. For comparison with previous rational features the heatmap axis is set similarly to Fig. 3B.

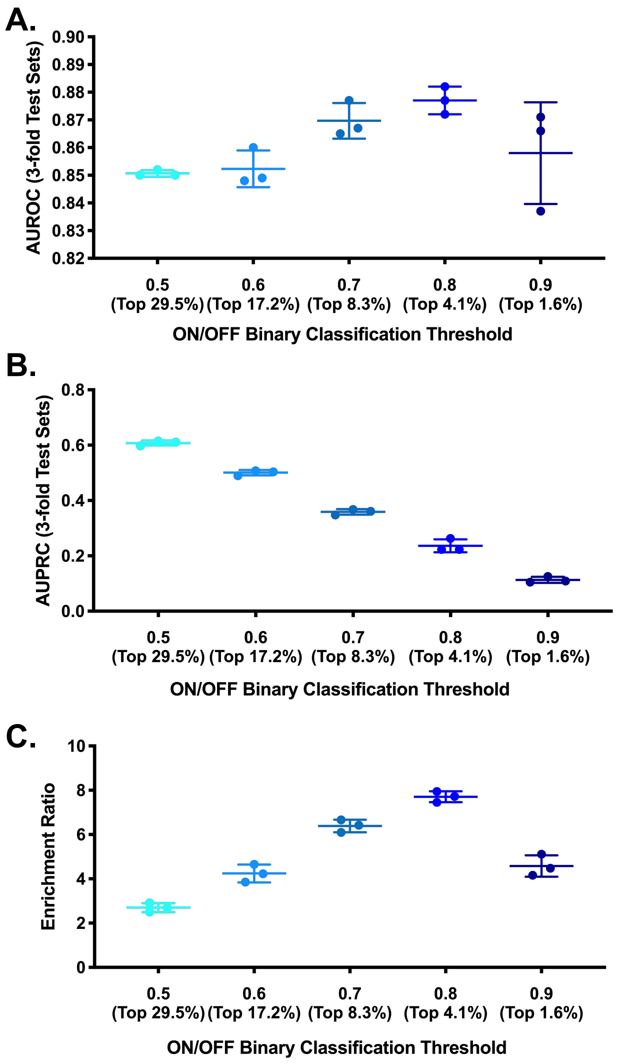

**Fig. S8. Determination of the optimal ON/OFF binary classification cutoff threshold.** AUC, P-R, and enrichment ratio analyses were used to determine the optimal cutoff threshold at which to binarize ON/OFF data for classification.

We trained a standard MLP architecture on the one-hot sequence representation of the toehold switch at five different binarization thresholds, and compared the following performance metrics: (A) model AUROC results, (B) model AUPRC results, and (C) model enrichment ratio over random chance. The enrichment ratio is calculated as the fraction of true positive toehold switches returned by the model (i.e., the precision) divided by the fraction returned by random chance. The enrichment ratio was specifically calculated at the level of precision for which the recall returns one positive switch per 100, or approximately ten on average for a typical mRNA of length ~1000nt. The final threshold selected for all classification models in this study was 0.7 (or the top 8.3% of switches), balancing a high enrichment ratio with a practical degree of overall precision.

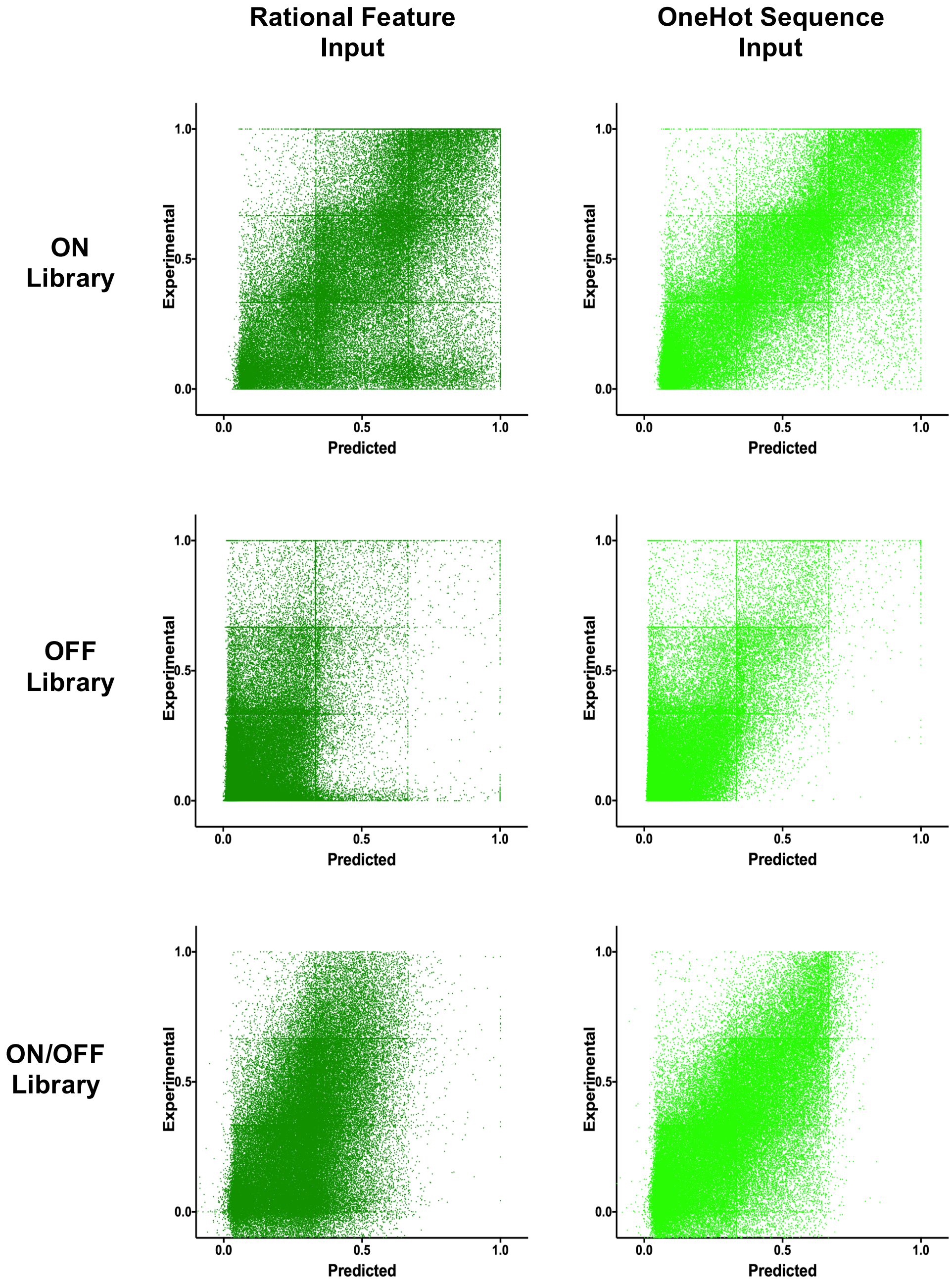

**Fig. S9. MLP predictions vs. experimental results.** Scatter plots of the predicted versus empirical values of our compiled test set are shown for ten-fold cross-validated MLP models trained with either the 30 pre-calculated rational thermodynamic features as inputs (left, dark green), or the toehold switch one-hot sequence representation as input (right, light green) for ON, OFF, and ON/OFF. Summary statistics are reported in Fig. 3D,E.

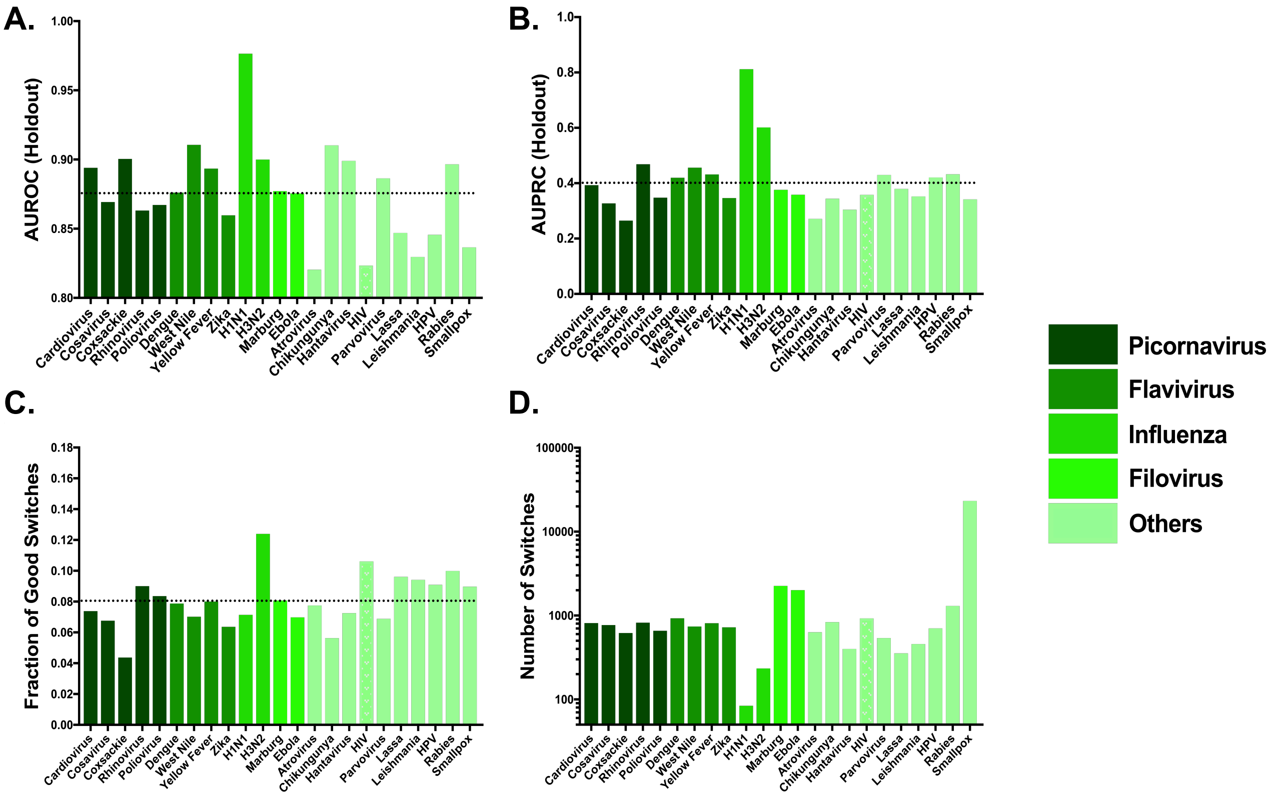

**Fig. S10. Holdout validation of individual viral genomes.** For each of the 23 pathogenic viruses tiled in our toehold switch dataset, every toehold switch targeting a given viral genome was withheld, and an MLP model was trained with the remaining sequences in the dataset using a one-hot sequence input representation classifying for ON/OFF ratio. The model performance was then evaluated on the switches of the withheld viral genoma as a test set. (A) Area under the receiver operating characteristic curves (AUROC) for holdout viral genomes. Dotted line denotes AUROC average across test samples. (B) Area under the precision-recall curves (AUPRC) for holdout viral genomes. Dotted line denotes AUPRC average across test samples. (C) Fraction of toehold switches in synthesized high-throughput library classified as high-performing for each virus type. Dotted line denotes average at 8%. (D) Total number of toehold switches synthesized for each virus type.

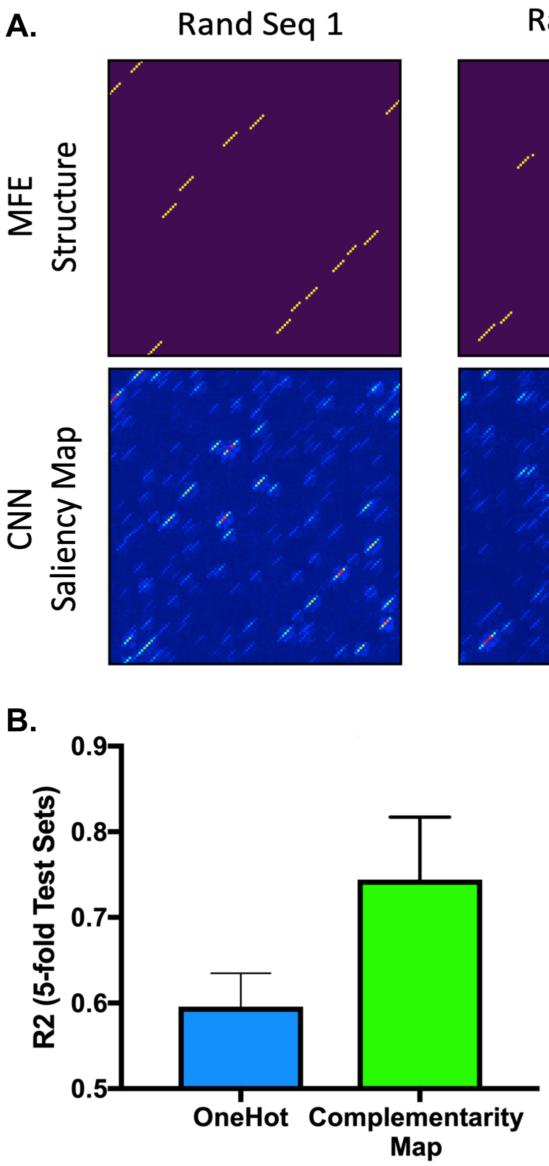

**Fig. S11. VIS4Map analysis of random toehold sequences in MFE predictor 2D CNN model.** A dataset of 50,000 random RNA sequences of length 120nt and their corresponding MFE values were generated using NUPACK. A convolutional neural network (CNN) was then trained to predict the MFE of each sequence using either a one-hot representation or a complementarity map representation of the sequence as input. (A) For three randomly selected RNA sequences, representative saliency maps generated from the CNN model are shown alongside the MFE structure pre-computed independently using NUPACK. The CNN model was trained on complementarity map inputs. Overlap between salient diagonal features in the VIS4Map outputs and MFE structure maps is visible. (B) We then compared the $R^2$ coefficients between NUPACK-calculated MFE values and the predictions of a CNN model trained either on a one-hot representation or a complementarity matrix representation of the random RNA sequences. Error bars show standard deviation from five shuffled test sets.

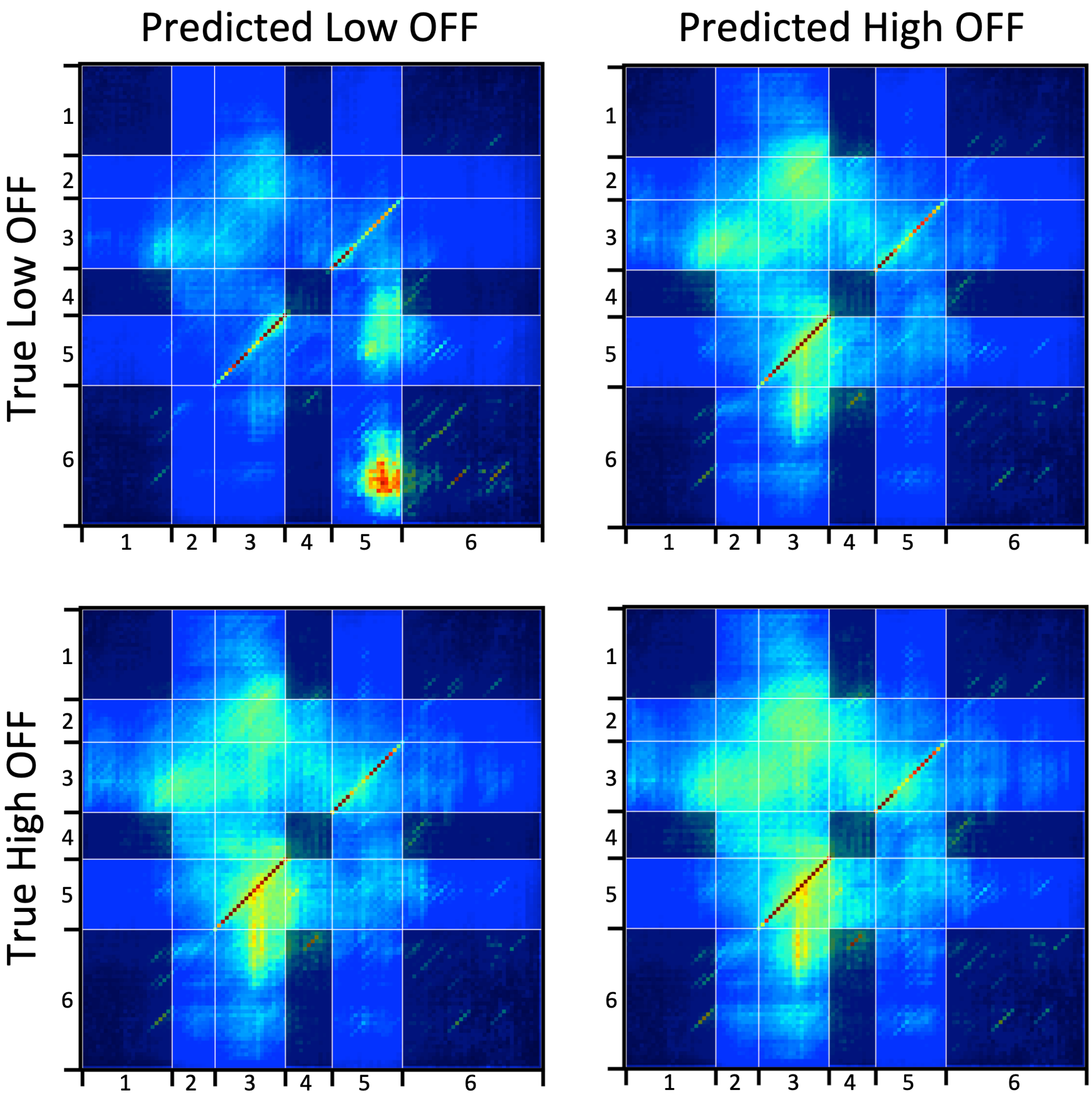

**Fig. S12. VIS4Map confusion matrix analysis of switch OFF conformation.** Saliency maps generated from a CNN model trained to predict the toehold switch OFF metric are shown for different ground-truth OFF metrics. The model was trained using a complementarity matrix representation of the toehold sequence as input. Regions labeled on the axes are as follows: 1) Constant Loop, 2) Toehold, 3) Ascending Stem, 4) Constant RBS Loop, 5) Descending Stem, and 6) Constant Linker. Regions of interaction between constant regions are shaded darker as they do not contain variability between different switch sequences. All saliency maps were generated from the test set only. Saliency maps were then sorted according to the 25% highest and 25% lowest experimentally-determined OFF signal. The 10% best-predicted and 10% worst-predicted saliency maps from the high OFF and low OFF groups were then averaged to produce the shown confusion matrix. Contrast was enhanced four-fold in the averaged maps in order to visualize more sparsely distributed features.

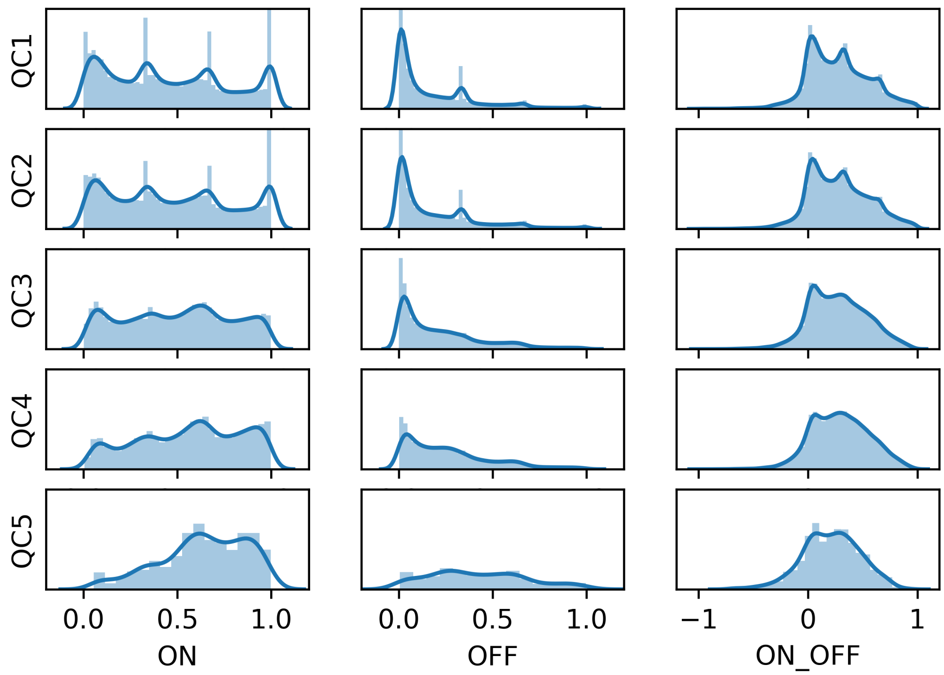

**Fig. S13. Dataset distribution vs. QC level.** **H**istograms of toehold switch library values for ON, OFF, and ON/OFF were grouped according to our five different QC threshold levels and are shown here for comparison. **The** y-axis limits are held constant for ON, OFF, and ON/OFF, respectively, across QC levels after normalizing for data subset size.

| | Quality Control Conditions | | | | Library Size | | |
|---|---|---|---|---|---|---|---|
| | OFF Count Threshold | ON Count Threshold | Upper Stdev. Cutoff | Lower Stdev. Cutoff | ON Variants | OFF Variants | ON/OFF Variants |
| QC1 | >= 5 | >= 5 | None | None | 126,620 | 180,552 | 110,931 |
| QC2 | >= 10 | >= 10 | None | None | 109,067 | 163,967 | 91,534 |
| QC3 | >= 20 | >= 40 | None | >0 | 77,040 | 90,264 | 43,044 |
| QC4 | >= 60 | >= 60 | 0.4> | >0.04 | 39,283 | 67,507 | 19,983 |
| QC5 | >= 300 | >= 300 | 0.4> | >0.04 | 6,187 | 12,551 | 1,137 |

**Table S1. Quality control thresholds.** The conditions for inclusion in our five quality control groups (QC1-5) are shown above, including standard deviation cutoffs and library count thresholds. QC2 was ultimately chosen as the final condition for inclusion in our dataset, and all data used or shown in this manuscript is for QC2 unless otherwise stated. The size of each dataset is shown in the three rightmost columns.

| | Library # | Trigger Sequence | On | Off |
|---|---|---|---|---|
| Low 1 | 1817 | CCGACACCTGTTTCATGGAACAATAAAAGA | 0.0153 | 0.0085 |
| Low 2 | 34792 | TGCTGTCTGTGAAACAGATAAATGGAAATA | 0.0176 | 0.0100 |
| Low 3 | 53587 | TCCCTTTCCCAGAAATAAACTTTTTTACCC | 0.0181 | 0.0136 |
| Low 4 | 72784 | TCACTGAGTCATTGCCATCTGCAGAATCAG | 0.0048 | 0.0134 |
| Low 5 | 104595 | TCCAAGACCCAAAGTTCTGGGAACTGGTGG | 0.0192 | 0.0156 |
| Low 6 | 158538 | TGGCAATTGTAGATATAACTTCTGGTAAAT | 0.0153 | 0.0183 |
| Low 7 | 188705 | ATCCAAATATAATGATGACCTATATGCCCT | 0.0158 | 0.0102 |
| Low 8 | 206071 | CCAATATGAGATCTGTAATGCTAACAGTTT | 0.0076 | 0.0146 |
| High 1 | 79874 | GTCATATAAAGGAAGAAGATAGGAGAAGAA | 0.9860 | 0.0031 |
| High 2 | 111242 | AGTTCACAAGAGATGGTTCATGGTGTTCCA | 0.9937 | 0.0132 |
| High 3 | 158916 | AAAGGTTAGCTTATGTTACATATCAAGATA | 0.9740 | 0.0016 |
| High 4 | 164714 | AATCACTGAAAATTGGAGTTAGGTATTGAC | 0.9747 | 0.0007 |
| High 5 | 166671 | GGTATGTTAAGTATGAGGCCTTATCCGTAC | 0.9895 | 0.0115 |
| High 6 | 187264 | TCAAGTTAGAGAAGGAAGTGGCTGAGACCC | 0.9856 | 0.0122 |
| High 7 | 215129 | TAAATCTATGAGAGATCAACGAAAAGGAAG | 0.9942 | 0.0150 |
| High 8 | 232933 | AAAGAAGAAATCATGCAAGAAAACAAAGGG | 0.9744 | 0.0007 |

**Table S2. Toehold switch sequences validated in cell-free format.** Sequences of the individually cloned toehold switches for cell-free validation using PURExpress were selected from the QC3 threshold. Their trigger sequences and flow-seq assay performances are shown (see Fig. 1F,S4 for cell-free assay performance). All highly-functional switches have ON/OFF of 0.97 or greater, while all poorly-functional switches have ON/OFF of 0.04 or less.

| ON Triggers | Motif | Counts in Foreground | Counts in Background | P-value | E-value |
|---|---|---|---|---|---|
| **Low versus High Signal** | | | | | |
| | UCUYUCU * | 349 | 0 | 7.10E-122 | 8.30E-117 |
| | GAUGG | 260 | 19 | 6.80E-63 | 7.90E-58 |
| | AAAAA | 391 | 128 | 1.90E-42 | 2.10E-37 |
| | CUCYUC * | 142 | 4 | 1.30E-39 | 1.40E-34 |
| | UAUUAAC | 123 | 0 | 1.70E-39 | 1.90E-34 |
| | UCUCAC * | 26 | 2 | 4.10E-37 | 4.50E-32 |
| | GAGUCGU | 100 | 0 | 5.80E-32 | 6.30E-27 |
| | GUUUUAUC | 100 | 2 | 8.50E-29 | 9.10E-24 |
| **High versus Low Signal** | | | | | |
| | ANSA | 785 | 427 | 6.00E-62 | 1.00E-56 |
| | AWUB | 644 | 359 | 9.50E-38 | 7.80E-33 |
| | UAYR | 355 | 163 | 3.90E-23 | 1.70E-18 |
| | GVRA | 270 | 128 | 8.20E-16 | 2.50E-11 |
| | ACK | 344 | 224 | 1.60E-09 | 3.80E-05 |
| | AUAA | 104 | 47 | 8.30E-07 | 1.40E-02 |
| **OFF Triggers** | **Motif** | **Counts in Foreground** | **Counts in Background** | **P-value** | **E-value** |
| **Low versus High Signal** | | | | | |
| | CNG | 762 | 503 | 8.40E-34 | 1.50E-28 |
| | GRS | 510 | 342 | 1.90E-14 | 1.80E-09 |
| | CCUH | 218 | 132 | 2.60E-07 | 1.60E-02 |
| **High versus Low Signal** | | | | | |
| | AWWWU | 591 | 346 | 2.10E-28 | 3.60E-23 |
| | WUAW | 472 | 333 | 1.40E-10 | 1.60E-05 |
| | AAAARA | 67 | 22 | 5.60E-07 | 4.30E-02 |

**Table S3. K-mer search results.** K-mer motifs searched with DREME using the trigger RNA sequences of the highest and lowest performing 1000 switches sorted by either ON or OFF signal. For this search, QC3 dataset was selected. * Denotes potential anti-SD pyrimidine-rich sequences.

| Rational Feature Sub-sequence Name | Sequence Region | Brief Description |
|---|---|---|
| SwitchOFF | 30-108 | Toehold switch off conformation |
| SwitchOFF-GFP | 30-144 | Off conformation with added GFP sequence |
| SwitchOFF-NoTo | 62-144 | Off conformation with toehold removed |
| SwitchON | 0-108 | Toehold switch on conformation |
| SwitchON-GFP | 0-144 | On conformation with added GFP sequence |
| Trigger | 0-29 | Trigger sequence alone |
| ToeholdOFF | 30-62 | Toehold region of switch including link1 |
| ToeholdON | 0-62 | Toehold region only hybridized to trigger |
| Stem | 62-108 | Stem only of toehold switch |
| AscendingStem | 62-100 | Ascending arm of the switch stem |
| DescendingStem | 80-108 | Descending arm of the switch stem |
| StemTop | 74-97 | Top half of the stem from start codon up |
| RBS-Linker | 80-134 | Region from RBS loop2 to linker |
| RBS-GFP | 80-144 | RBS-Linker with added GFP sequence |

| **[-3,-1]** | **[0,29]** | **[30,49]** | **[50,79]** | **[80,90]** | **[91,96]** | **[97,99]** | **[100,108]** | **[109,134]** | **[135,144]** |
|---|---|---|---|---|---|---|---|---|---|
| GGG | trigger | loop1 | switch | loop2 | stem1 | AUG | stem2 | linker | post-linker |

**Table S4. Rational feature sub-sequences.** The sub-sequences from which the thirty rational features used as MLP input were calculated using ViennaRNA are shown here in the upper panel. In the lower panel, we show the full un-truncated toehold switch sequence framework from which the sub-sequences in the top table were selected.

### Supplemental References

1. A. A. Green, P. A. Silver, J. J. Collins, P. Yin, Toehold switches: de-novo-designed regulators of gene expression. *Cell* **159**, 925-939 (2014).
2. K. Pardee *et al.*, Rapid, low-cost detection of Zika virus using programmable biomolecular components. *Cell* **165**, 1255-1266 (2016).
3. K. Pardee *et al.*, Paper-based synthetic gene networks. *Cell* **159**, 940-954 (2014).
4. S. E. Hunt *et al.*, Ensembl variation resources. *Database* **2018**, (2018).
5. P. Oberacker *et al.*, Bio-On-Magnetic-Beads (BOMB): Open platform for high-throughput nucleic acid extraction and manipulation. *PLoS biology* **17**, e3000107 (2019).
6. A. Espah Borujeni, H. M. Salis, Translation initiation is controlled by RNA folding kinetics via a ribosome drafting mechanism. *Journal of the American Chemical Society* **138**, 7016-7023 (2016).
7. T. L. Bailey, DREME: motif discovery in transcription factor ChIP-seq data. *Bioinformatics* **27**, 1653-1659 (2011).
